# *A Priori* recognition for biological phenomena prior to empirical understanding

**DOI:** 10.1101/2025.10.02.680169

**Authors:** Yuji Takeda, Junji Yokozawa, Risako Yamaguchi, Shinichi Saitoh, Hironobu Asao

## Abstract

Biological phenomena include unrecognized events. These unrecognized events often unknowingly increase observer bias, which inhibits open and reproducible science. In this study, we focused on the recognition procedures underlying primary data and modeled cell population behavior. Using agent-based modeling (ABM), we demonstrated that cellular behaviors can be categorized into 11 distinct types, a framework we defined as the Behavioral Eleven Cell Class (BECC). We further validated BECC by describing the differential STAT3 phosphorylation patterns in leukocyte subsets and polarized T cell differentiation in OT-II transgenic mice. BECC serves as a novel descriptive method with three defining features: (i) it functions as a symbolic system independent of numerical or linguistic constraints, (ii) it represents the minimal unit of recognition, and (iii) it enables the expression of inherently unrecognizable phenomena. BECC allows for observer subjectivity and the relativity of results while enforcing rigorous recognition. This unique approach provides a practical and conceptual basis for advancing open and reproducible science.

## Introduction

The observational results of biological phenomena may include unexpected and unrecognizable events (Hicks 2023, Hunter 2017). Therefore, even under identical conditions, observations conducted at different times and places may yield different results. Moreover, even when the numerical values differ, they may still reflect observations of the same phenomenon. Utilizing observational results obtained at different times and locations is not straightforward. This can be regarded as a reproducibility issue caused by an unrecognizable event.

To understand biological phenomena, it is necessary to describe the observed results using language in manuscript. However, language allows for diverse expressions and evolves over time, so the same phenomenon may be described differently. This is a problem of linguistic fluctuation (i.e., arbitrariness and flexibility in language). Furthermore, exaggerated understandings often result in misperceptions (Chiu et al. 2017). This issue can be described as a problem of linguistic fluctuation.

The process of recognition shapes our understanding of phenomena (see Premise 1). A different recognition of a concept or phenomenon can lead to an updated understanding and generate new insights. However, the communication of new recognition is entrusted to citation of paper. This citation activity, however, tends to create a self-reinforcing cycle in which papers that are easier to cite are more frequently cited (Ross-Hellauer et al. 2022). This cycle can be problematic. For observer (see Premise 2), the statistical methods, databases, algorithms used for analysis, and the understanding derived from dualistic interpretations may influence and constrain different recognition. This issue can be framed as a problem of conservative recognition.

Issues of reproducibility, linguistic fluctuation, and conservative recognition are not sufficiently resolved by improvements to institutions, collective knowledge, and search algorithms alone. Therefore, we focused on and attempted to improve the methods for recognizing biological phenomena themselves. Accordingly, this study proposes the “Behavioral Eleven Cell Class (BECC)” as a new method for recognizing biological phenomena. This classification method has three characteristics: 1) it is symbols distinct from language, converging on a single meaning without generating new meanings through combination, similar to map symbols, pictograms, or fixed signs such as +,-, and =, 2) it can express the presence of unrecognizable phenomena, and 3) it can express the minimal unit of recognition (see Premise 3). These three characteristics provide a means of addressing the problems of reproducibility, linguistic fluctuation, and conservative recognition. To demonstrate this potential, we verified that BECC could be universally applied to biological phenomena using an abstract model based on an agent-based modeling (ABM) and proved the validity of its logical background. As an example of BECC, we describe the activation of cellular signaling pathways, and use OT-II mice, which are widely used in immunology, to describe T cell differentiation.

## Materials and methods

### Blood collection from healthy volunteers

This study was approved by the Ethics Committee of Yamagata University Faculty of Medicine (approval number: 2024-44). Peripheral blood was collected from healthy volunteers (male, 37 to 55 years of age) after obtaining written informed consent and anticoagulated with 5 U/mL of low-molecular-weight heparin. Blood collection was performed on different days for each person and treated as independent experiments.

### OT-II transgenic mice (OT-II)

OT-II mice express T-cell receptor (TCR) α-and β-chains that recognize the MHC class II Ib-restricted OVA peptide (residues 323–339) in the C57BL/6J background. They were kindly provided by Dr. W. Heath (WEHI, Melbourne, Australia). The animal experiments were approved by the Animal Experiment Committee of Yamagata University Faculty of Medicine (approval number, R7015). The mice were bred at the animal facilities of Yamagata University Faculty of Medicine, under specific-pathogen-free conditions and were used for tail sampling between 5–6 weeks of age.

### ABM simulation

ABM simulation was performed using NetLogo program (https://www.netlogo.org/) and the file (Model of cell population version 5.0.2). The parameters for each simulation are described in Figure legends. The description of the interface screen, parameter manipulation, and agent behavior are provided in Fig. 2b and Supplementary Information. The code of “Model of cell population version 5.0.2” is also available in the Supplementary file.

**Fig. 1.**
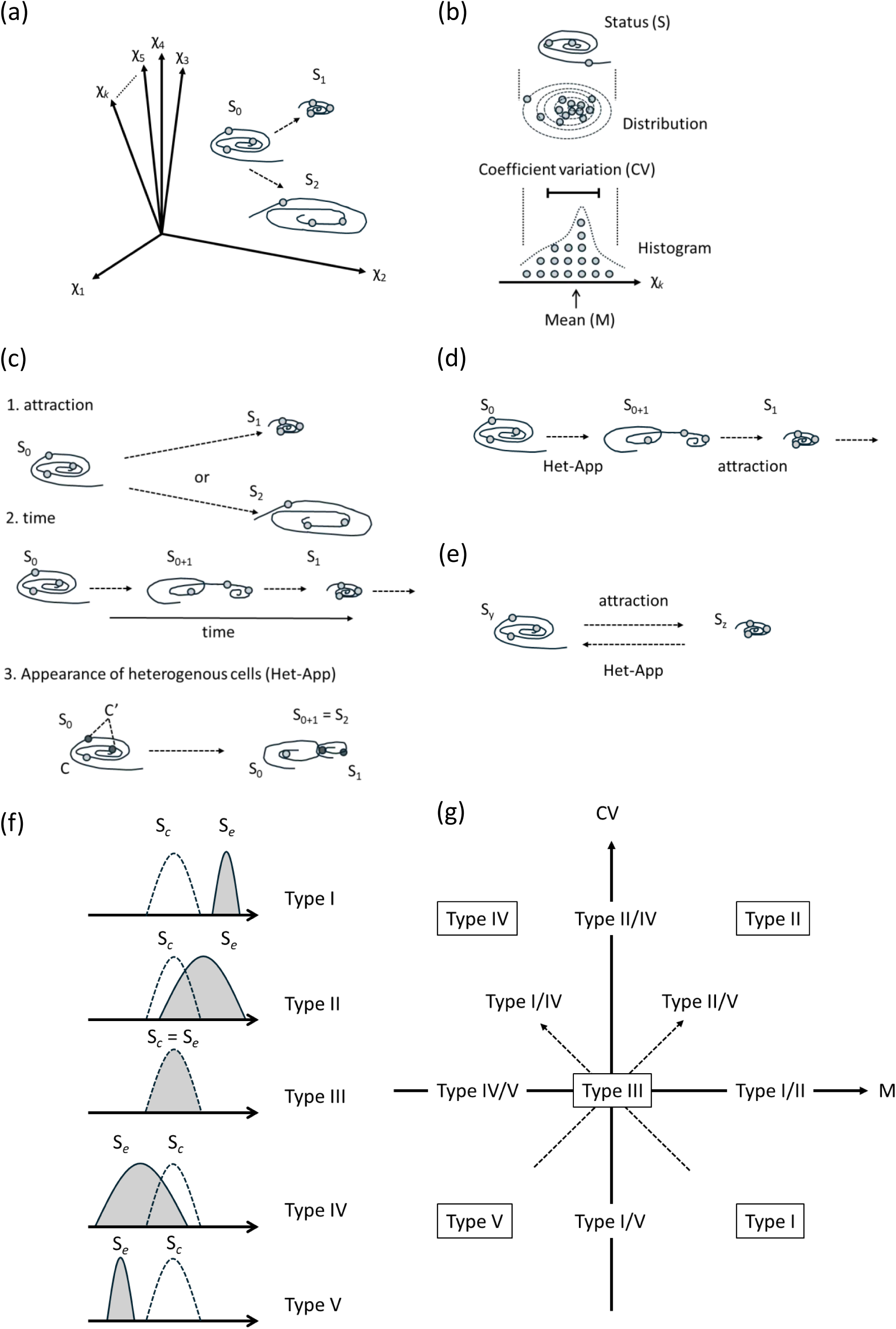
Schematic representation of relationship between cell population status and cell population distribution. The gray circles represent a representative example of a single cell, and the solid line, like a vortex, represents the trajectory of the cell behavior over a fixed elapsed time. The dashed arrows indicate the direction of cellular change after stimulation. (a) Cellular responses in multi-dimensional space. A cell responds from initial state S to reacted state (S1 or S2) upon stimulation, showing cellular variation in *k*-dimensional space. The *k*-dimension does not include the time axis. (b) Distribution of similar cell states and cell populations. When a collection of similar cells is observed, the cell state can be detected as a distribution of cell populations (dashed ellipses). A distribution of cell populations can be measured as a histogram for any parameter. The state of the cell population, measured as a histogram, can be informed as a mean value (M) and a coefficient of variation (CV). (c) Factors that change the distribution of cell populations. The factors that change the distribution of the cell population include 1. attraction, 2. time, and 3. appearance of heterogeneous cells (Het-App). The cells in C’, indicated by the black circle, are heterogeneous cells compared to the cells in C, indicated by the gray circle. Heterogeneous cells are unrecognizable and inseparable in *k*-dimensional space. (d) The time factor consists of Het-App and attraction. (e) The increase or decrease in the distribution of a cell population is an inducement and the appearance of heterogeneous cells. Comparing cell state “y” and cell state “z”, the increase or decrease in the distribution of the cell population is related to attraction and Het-App, respectively. (f) Five classifications based on histogram comparisons. The five patterns that can be recognized when comparing the histogram of Sy (dashed, unfilled) for the control group of a cell population and the histogram of Sz (solid, filled in gray) for the experimental group. (g) The Behavioral Eleven Cell Class (BECC) of these 11 categories is represented by a matrix with two axes (solid arrows) for CV and M. The dashed arrows indicating Type I/IV and Type II/V show the negative and positive correlation of CV and M, respectively.

**Fig. 2.**
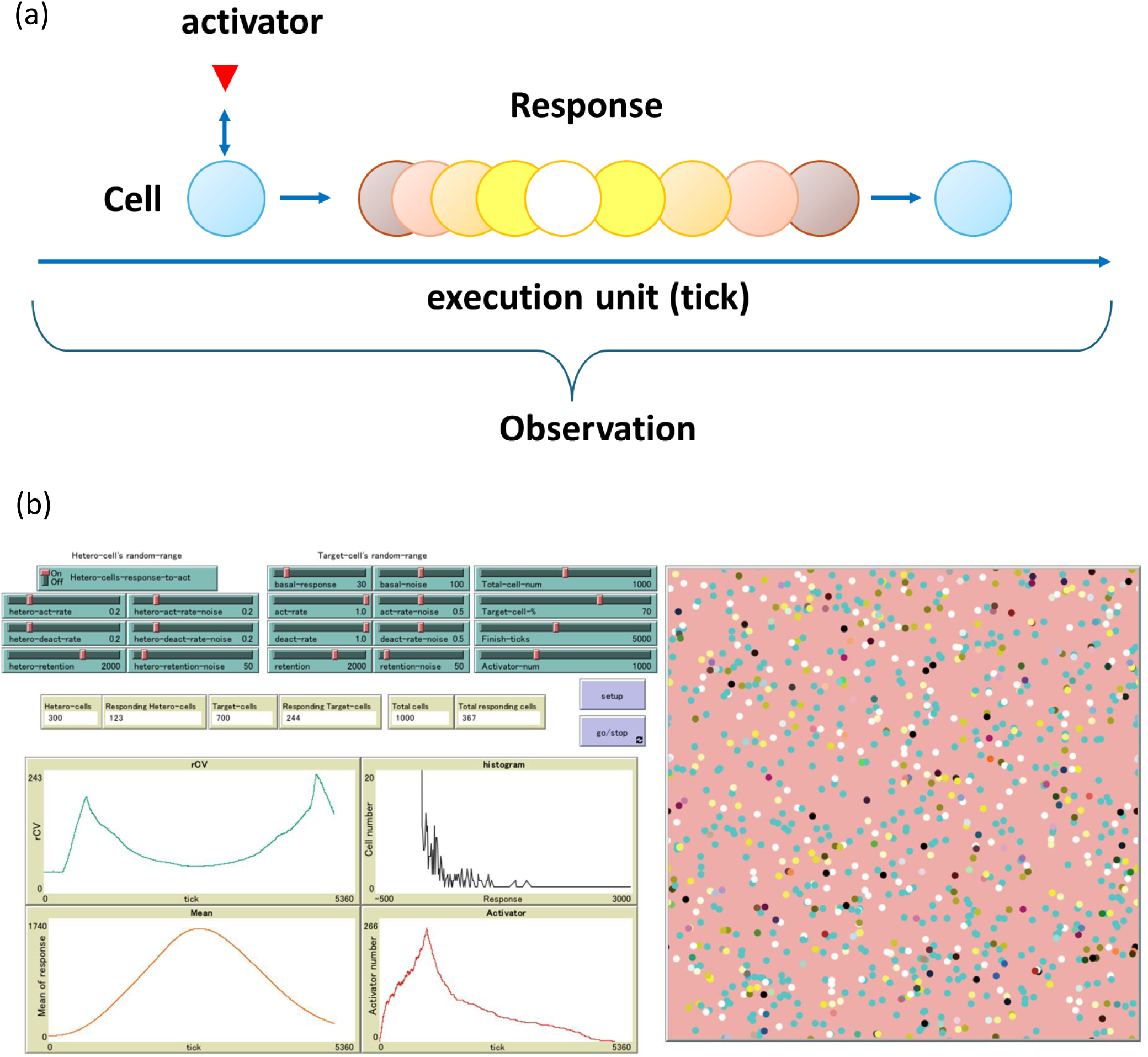
Modeling biological responses using an agent-based model. (a) Representation of biological responses by agents. A “cell” is represented by a circle (○). A stimulation is represented by the encounter of an “activator” (△) with a cell (○). A “response” is expressed as a response value (in the figure, a change in color tone). “Observation” is obtained as a numerical response value, which is then represented by the mean value (M) and the coefficient of variation (CV) derived from the histogram of response values. Any elapsed time is represented by a single tick for each agent to act once, which is done according to the program code. (b) The interface screen of the Model of cell population_ver.5.0.2 file created by NetLogo. A detailed description of each panel comprising the interface screen is provided in Supplementary Information.

### Human blood stimulation with cytokines

Following sample collection, 0.1 mL of blood per tube was transferred into 1.5-mL microtubes and pre-incubated for 60 min at 37℃. After pre-incubation, samples were mixed with vehicle (Roswell Park Memorial Institute (RPMI) 1640 medium), Interleukin-21 (IL-21) (10 nM; PeproTech, Rocky Hill, NJ, USA), or G-CSF (10 nM; Chugai Pharmaceutical, Tokyo, Japan) and stimulated for 15 min at 37℃. Stimulated blood was immediately fixed in 1.4 mL of 1x BD Phosflow^TM^ Lyse/Fix buffer (Becton, Dickinson and Company (BD) Biosciences, San Jose, CA) for 10 min at 37°C. The fixed cells were pelleted by centrifugation at 800 × *g* for 1 min at 24°C and then suspended in 90% methanol (0.3 mL/tube) at −20°C until staining with antibodies.

### Human blood cell staining for flow cytometry

The fixed cells in methanol were washed twice with 0.8 mL phosphate-buffered saline (PBS) and suspended in PBS containing 3% fetal calf serum and 0.1% sodium azide. The cells were incubated with antibodies, including Pacific Blue anti-human CD3 monoclonal antibody (mAb; clone UCHT1), allophycocyanin-conjugated anti-human CD16 mAb (clone 3G8), and Alexa488-conjugated anti-pY705-STAT3 mAb (4/P-STAT3), for 30 min at 22–25°C after fragment crystallizable receptor blocking using fragment crystallizable blocker (Human TruStain FcX, Biolegend, San Diego, CA, USA). After the reaction, the cells were washed with PBS and analyzed by flow cytometry (FACSMelody, BD Biosciences). Cell debris and doublets were excluded from the analysis by forward-and side-scatter gating. Mean fluorescence intensity and robust CV were analyzed using FlowJo software (version 10.10.0, BD Biosciences).

### Mouse T-cell staining for flow cytometry

The tip of the mouse tail was cut, and approximately 0.1–0.2 mL of peripheral blood was collected in microtubes containing 5 mM ethylenediaminetetraacetic acid-added PBS. After centrifugation and removal of the supernatant, the blood cells were stained with antibodies including fluorescein isothiocyanate anti-mouse CD4 mAb (clone RM4-5), Pacific Blue anti-mouse CD3 mAb (17A2), Phycoerythrin anti-mouse TCR Vα2 mAb (B20.1), and/or PE anti-mouse CD8α mAb (53-6.7), at 4°C for 30 min. After the reaction, the blood cells were fixed in 1x BD PhosflowTM Lyse/Fix buffer and then washed with PBS. The washed cells were analyzed by flow cytometry (FACSMelody), as described above.

### Statistical analysis

The statistical methods, sample sizes (n), and *p* values for each experiment are described in the Figure legends. Statistical analyses were performed using Prism software (version 5.03, GraphPad Software, San Diego, CA, USA). A *p* value of *p* < 0.05 was considered statistically significant.

## Premise in this study

### Premise 1: Perception, recognition, and understanding

The process of human understanding, as described in cognitive psychology textbooks, begins with perception (using the five senses) to observe the world. This is followed by recognition, where we pay attention to and select meaningful information from those observations. Finally, we achieve understanding by constructing and interpreting the meaning of what we have recognized.

This paper is premised on the idea that the understanding of biological phenomena follows a similar process. The resulting understanding is then documented in writing. In science, our perception is, of course, now being extended by experimental equipment and information processing technology. For the purpose of this paper, “recognition” specifically denotes the intermediate step in this process that leads to understanding.

### Premise 2: Affirmation of the observer’s existence

Scientific facts are based on objective evidence, not subjective experience. Therefore, a statement like, “We saw what appeared to be an increase in X,” is considered weak. A scientific claim must be made with certainty, such as, “There was a measurable increase in X.” However, the complete removal of the subjective observer from the scientific process has become a mere formality.

Human analysis of the human brain is not subject to criticism for being a cognitive cycle. This is because the observer observes and measures the brain as “brain of not human but another person”. This is an example of the classic observer’s affirmation. This observer’s affirmation is not limited to brain science.

Our understanding of many biological phenomena has advanced rapidly. However, this progress has also led to a proliferation of “unrecognizable differences.” For example, even if the specificity of ligands and receptors is proven, their signals are still interpreted through universal model of signal-transduction pathways. In other words, although the binding is specific, it can still be represented as universal phenomena. Therefore, it is assumed that there are “unrecognizable differences” in the signal transduction pathway. When we try to understand that these “unrecognizable differences” consist of various combinations, such as different co-stimuli, variations in transcription factor patterns, stimulus strength, etc., we must prepare that new mechanisms are added to enable the classification of various combinations. Consequently, new understanding constitutes a new “unrecognizable difference,” much like a fractal structure. Alternatively, if the accumulation of understanding is likened to the volume of a sphere, then as understanding increases, so does the implicit, unrecognizable differences, similar to the surface area of the sphere.

To unconsciously accept and share these inherent limitations—which we have been calling “unrecognizable differences”—as a valid basis for our observations, or subtle variations or entirely new phenomena that are not yet apparent to us, we employ a form of “limited recognition”. This approach is based on the premise that we are limited observers, and that the biological event we observe is merely a specific subset of all possible observations, as defined by the observer’s tools and methodology. Without accepting these limitations, we could never arrive at an understanding due to the overwhelming number of variables. When we accept that is a tentative understanding, we already allow for the existence of an observer who has declared a specific range of observations.

### Premise 3: To recognize biological phenomena, compared to a control group

A distinguishing characteristic of biological science is that we assume we are observing a complex system. For a finding to be considered a meaningful recognition, any observed difference must be shown to have a potential effect on the entire complex system. We achieve this recognition by comparing the subject to a control group, which we assume is similarly complex.

For example, a reaction process that is fully predictable by computation is not considered biological recognition. The prediction becomes a recognition only when it is accompanied by an explainable meaning that puts the process into a larger biological context. This is similar to how a weather forecast becomes meaningful recognition only when we understand the social consequences of rain versus no rain.

Another example can be seen in the classification of morphological characteristics. A simple description of a new organism’s location and physical traits is a factual report of the discovery, but a comparison to previously known species is essential for it to be recognized as a new finding. This process of identifying and explaining differences through comparison is the key recognition that forms the basis for a deeper understanding of biological phenomena.

This structure of recognizing through comparison is relative, making the selection of the control group arbitrary. However, the recognition gained from the comparison of these two entities serves as the minimal unit of recognition for understanding.

## Confirmation in this study

### Confirmation 1: Time

The precise definition of “time” is left to the fields of philosophy and physics, as it concerns the very nature of its existence. In this paper, we consider the recognition of a biological phenomenon to begin with an observer’s act of observation. Therefore, the concept of “time” here also includes the observer’s perception. In this context, there may be no unified time between the observed object (e.g., a cell) and the observer (see Confirmation 6). Thus, “time” in this paper refers to phenomena that an observer assumes have changed in an irreversible, unidirectional manner.

### Confirmation 2: Observation results are detached from time

When an observer observes a biological phenomenon, they capture a specific moment and quantify the measured variables. Even with morphological observations or time-lapse studies, the results are quantified and compared over a set period. Ultimately, observing a biological phenomenon becomes an act of comparing “photographs” (or matrices of numerical data) taken with a fixed exposure time. In doing so, the observed or calculated results are detached from the irreversible, unidirectional flow of time.

### Confirmation 3: Reason for using the coefficient of variation (CV) instead of standard deviation (SD)

When measuring the same parameter across different spatiotemporal conditions, the measurement methods or instruments used may have different units or scales. Because standard deviation (SD) assumes a normal distribution, comparisons using SD require the values to be on the same unit and scale.

This paper, however, assumes scenarios where measurement results are compared under varying spatiotemporal conditions. Therefore, when comparing values with different units or scales, the coefficient of variation (CV) is more appropriate.

Furthermore, the robust CV is even more suitable. It is based on the median rather than the mean, making it less affected by sample size and free from the assumption of a normal distribution. For the remainder of this paper, all references to CV refer to the robust CV.

### Confirmation 4: Differences between ideal responses and cell populations

When an observer examines cellular responses to a stimulus, they compare a group of cells that they have arbitrarily collected and designated as similar. If an observer assumes an ideal response exists, measurements are interpreted as if they were taken from a uniform, ideal cell population, and the actual distribution of the cells is treated as experimental error. Consequently, assuming an ideal response makes it impossible to classify using BECC or to identify the responses of individual cells.

It is important to note that certain methods—such as Western blot, qRT-PCR, bulk RNA-seq, and binary event-based assays (e.g., positive vs. negative)—involve processes that presume a uniform cellular response by lysing the cells before measuring any cellular heterogeneity. Therefore, even if four replicate measurements are performed for statistical analysis, the resulting distribution only reflects the experimental error from measuring four instances of a single, equivalent cell.

In contrast, BECC, as proposed in this paper, assumes an experimental system in which cell populations are measured. An appropriate number of cells (more than 100, which is a sufficient number to ensure that percentile calculations are not affected by differences in formulas) are included to derive a valid CV. Currently, measurement techniques such as flow cytometry, which independently assess individual cells, are useful for BECC.

### Confirmation 5: Comparing different conditions and temporal changes on ABM simulation

The minimal unit of recognition in observational results is defined as the difference between a control group and an experimental group. These differences can be categorized into two comparison methods:

- Comparing different conditions at the same observation point.
- Comparing different observation times under the same condition.

Naturally, it is also possible to compare different observation points under different conditions, which involves analyzing the interaction between “time” and “different conditions.” This is a statistically valid form of comparison. However, since the focus of this study is on identifying the minimal unit of recognition, we will not conduct comparisons that involve combinations of more than two different conditions.

In this study, the temporal changes of agents representing biological phenomena are tracked as a history of updates that occur at each tick. Here, a tick represents one execution of a behavior by each agent according to the program code (the execution unit). Based on this framework, we can draw comparisons using two methods:

- The difference in CV and Mean (M) at a single tick (observation point) between two different simulation conditions.
- The difference in CV and M between two different ticks (observation points) within the same simulation condition.

These two methods form the basis for our experimental design.

### Confirmation 6: In ABM simulations, a “tick” is not a measure of time, but a unit of execution

At first, this section considers how an experimental group can yield the observation that no progressive change has occurred. If neither the experimental nor the control group changes, it is impossible to recognize time as progression. Therefore, the observer must rely on an external factor that changes progressively—what we call a “clock.” Without a clock, it becomes impossible to observe that “no change has occurred.” However, introducing a clock to the control group in such a scenario would violate the rule of minimal recognition units.

In the ABM simulation discussed here, we do not use a clock based on quantum oscillations. Instead, a “tick”—where each agent performs one action based on the program code—serves as a single unit of execution for a biological phenomenon. This unit of execution does not necessarily correspond to quantum oscillations. In this study, if the unit of execution of biological phenomena is constant, then quantum oscillations can be considered as fluctuating.

This observational method allows us to determine that’no change has occurred’ even without a clock, making it a minimal unit for recognizing change (= minimal unit of recognition). A key point is that if the observer’s unit of biological execution differs from that of the observed cells, there is no common unit of execution. This difference creates an observer-induced error across experiments. Thus, when an observer uses a clock to measure a cell’s response, the cell’s unit of execution will not be constant and will fluctuate.

### Confirmation 7: In ABM simulations, two types of errors can be separated and handled independently

In scientific research, when measurement accuracy is in doubt, repeating experiments can increase statistical power and compensate for errors originating from the observer. The statistical power of detection ultimately determines BECC classification, meaning the number of repetitions and the type of statistical analysis are critical components of the observer’s perspective.

In ABM simulations of biological phenomena, these two types of errors can be separated and handled independently. For example, in the Cell Population ver. 5.0.2 model, we set a range of random values (noise) for each parameter. This noise represents the inherent error from biological fluctuations.

On the other hand, observer error occurs when the observer cannot precisely determine how many ticks (execution units) have passed. In other words, this reflects a state where the execution units of the observed target and the observer are not aligned (see Confirmation 6). Specifically, the observer’s experimental error is represented not by the distribution of response values collected at a single tick, but by the distribution of response values collected over a range of ticks.

When the error from biological fluctuations is equal to or greater than the observer’s error, a statistically significant difference will be detected if one truly exists. Conversely, if no significant difference exists, increasing the number of repetitions will not yield a significant result. Therefore, it is unnecessary to assume scenarios where biological fluctuation error exceeds or equals observer error.

## Results

### Biological phenomena can be represented within multi-dimensional space, and it is reasonable to assume the existence of an ideal one-dimensional parameter for observing cell behavior

In recent years, comprehensive analyses of individual cells using techniques such as high-parameter flow cytometry and single-cell RNA sequencing (scRNA-seq) have advanced our understanding of various biological phenomena. As recognized through these comprehensive approaches, cellular behavior can be also represented within multi-dimensional space. However, sometimes a specific biological phenomenon is revealed through the upregulation of a single characteristic gene. Yet even then, numerous molecular processes, such as epigenetic regulation, transcription factor activation, and mRNA translation, contribute. Therefore, even without comprehensive analyses, it is possible to represent cellular behavior within multi-dimensional space. This suggests that a single measurement can reflect an observation derived from composite biological events as described previously (Takeda et al. 2022, Takeda et al. 2017). Based on the previous paper, we re-evaluated this framework by adding the concept of time and provided a novel explanation in this study.

If we attempt to represent cellular behavior as a series of snapshots within multi-dimensional space that does not include the time axis (see, Confirmation 1 and 2), it can be illustrated as shown in Fig. 1a cell in a given state (S) is confined within a specific region (Fig. 1a). When examined at the population level rather than a single cell, each individual cell is influenced independently by various factors while receiving distinct effects from specific factors. Consequently, the cell population can be observed as a distribution of cells occupying a region with certain probabilities. When this distribution is viewed through a single parameter, it can be visualized as a histogram. Naturally, multiple histogram perspectives exist in multi-dimensional space, each representing cell frequency by height. The choice of perspective depends on the observer’s selection of measurement parameters. The viewpoint derived from one such parameter is termed the “ideal one-dimensional parameter.” The coefficient of variation (CV) and mean (M) of the histogram along this ideal one-dimensional parameter reflect cellular behavior (Fig. 1b; see Confirmation 3). In summary, biological phenomena can be represented within multi-dimensional space, and an “ideal one-dimensional parameter” may allow cellular behavior to be observed.

One might question whether the assumption of an “ideal one-dimensional parameter” truly holds for multi-dimensional cellular behavior. When comprehensive multi-parameter analyses are conducted, multivariate techniques such as principal component analysis (PCA) or Uniform Manifold Approximation and Projection (UMAP) are often employed to reduce data dimensionality for characterizing cell populations. The statistically synthesized dimensional axes in these analyses essentially aim to approximate an ideal one-dimensional parameter that highlights the differences between groups. Even if the resulting synthetic dimension cannot be reduced to a single measurable item, it suggests that an ideal one-dimensional parameter may exist, effectively compressing multi-dimensional cellular behavior into fewer dimensions. Multiple ideal one-dimensional axes may exist, and in some cases the observer may not be able to identify any such parameter. In this study, the measurement parameter of interest to the observer was treated as an ideal one-dimensional parameter. This ideal unidimensional item is the opposite of the null hypothesis (assuming no difference) and is a measurement item that is assumed to reflect cell behavior.

### The comparison along a one-dimensional parameter represents the minimal unit of recognition

The recognition of biological phenomena is thought to arise from comparisons with a control group (see Premise 3). In this context, comparison along a one-dimensional parameter represents the “minimal unit of recognition” for observers. This is because quantitative information from a single parameter is represented as a histogram. Comparing this histogram with that of the control group constitutes the most basic form of comparison. Just as the shortest distance between two points is a straight line, a fundamental axiom, comparing two elements is the simplest and most intuitive form of analysis.

Typically, attention is focused solely on the statistical significance of increases or decreases in measured values, whereas the CV, which reflects the distribution of values, is often overlooked. However, when considering cellular behavior within multi-dimensional space, the distribution of cells can provide valuable information. In the simplest comparison procedure, which involves comparing two histograms, the maximum information that can be extracted consists of two aspects: changes in the measured values and changes in the CV. These two elements form the “minimal unit of recognition” for detecting meaningful differences.

### CV changes due to “Attraction” or “Appearance of heterogeneous cells (Het-App)”

It is common practice to treat the mean values of the measurement parameters as useful information. CV, on the other hand, has not received much attention. The usefulness of CV is illustrated through the following thought experiment. When a cell responds to a stimulus, multiple observable parameters may change. If we represent these changes within multi-dimensional space, they appear as a shift in the distribution of cells from one state (S_0_) to another state (S_1_ or S_2_). In this context, the distributional changes arise from three possible conditions (Fig. 1c).

(1) Attraction: Attraction is defined as a set of forces acting to keep cells in a particular state. When a cell population transitions to state S_1_, in which the distribution is narrower than state S_0_ owing to stimulation, it indicates an increase in attraction compared to state S_0_. Conversely, if stimulation produces a more dispersed state S_2_, then S_2_ is considered to have decreased attraction compared to S_0_.
(2) Time: During the transition of a cell population from S_0_ to S_1_, cells exist in both states simultaneously. At this point, the cells are captured in a state represented as S_0_ + S_1_, which is the fusion of both distributions. Additionally, there may be a period following the completion of the transition during which the overall distribution of cells narrows.
(3) Appearance of heterogeneous cells (Het-App): The initial cell population C in state S_0_ exists, and a heterogeneous subpopulation C’ emerges, indistinguishable from C under current multi-dimensional parameters. C and C’ cannot be clearly separated along any ideal synthetic axis. If only C’ transitions to state S_1_, then both C in S_0_ and C’ in S_1_ coexist. Therefore, C, which includes C’, is captured in the fused state S_0_ + S_1_, resulting in an increase in the overall cell distribution.

The increase or decrease in CV caused by “time” is observed when observers compare two points along the time axis. For example, when comparing “pre-stimulus” and “post-stimulus,” the two groups “before” and “after” are unconsciously integrated within a framework of continuous change along an irreversible direction and recognized as a continuous change (see Confirmation 1 and 2). If we exclude the integrating observational perspective of “time,” the situation reduces to comparing two points. Thus, the increase or decrease in CV caused by “time” consists of two components: “attraction” and “Het-App” (Fig. 1d).

When CV decreases, even if heterogeneous cells are present, their response leads to a smaller distribution, meaning heterogeneous cells are effectively masked by “Attraction.” Therefore, when CV decreases, it is unnecessary to refer to the “presence of heterogeneous cells” in the control group (Fig. 1e). In contrast, when CV increases, it indicates that within a cell population normally constrained by “Attraction,” easily deviating heterogeneous cells emerge (Het-App). Therefore, when CV increases, it is unnecessary to refer to the “attraction” state of the control group (Fig. 1e). In summary, when CV decreases, it is integrated as the occurrence of “Attraction” along the ideal one-dimensional parameter. Conversely, when CV increases, it is integrated as the emergence of “heterogeneous cells” along the ideal one-dimensional parameter.

### The minimal unit of recognition is composed of M and CV, and cell population behavior can be classified into 11 types based on the dynamics of M and CV

The control state of the cell population (subscript-italic *c*), S*c,* is represented by the histogram of CV*c* and M*c*, and the examined state of the cell population (subscript-italic *e*), S*e,* is represented by the histogram of CV*e* and M*e*. When the histogram of S*e* is compared with that of S*c*, an increase in M is represented by a rightward shift, and a decrease in M is represented by a leftward shift. Considering this in combination with increases or decreases in CV, the patterns of histogram changes can be categorized into five types (Fig. 1f). Based on previous studies examining the activation of signal transduction pathways, these five patterns can be described as follows:

1. M*c* < M*e* and CV*c* > CV*e* (M↑, CV↓) is defined as Type I, Attractive.
2. M*c* < M*e* and CV*c* < CV*e* (M↑, CV↑) is defined as Type II, Subsequent.
3. M*c* = M*e* and CV*c* = CV*e* (M→, CV→) is defined as Type III, Passive.
4. M*c* > M*e* and CV*c* < CV*e* (M↓, CV↑) is defined as Type IV, Counter.
5. M*c* > M*e* and CV*c* > CV*e* (M↓, CV↓) is defined as Type V, Negative arbiter.

The linguistic representation of the pseudonyms is provided as a means of facilitating analogy with the cellular behavior represented by the symbols Type I through Type V.

Questions may arise regarding the interpretation of distorted histograms or those with multiple peaks. Distorted histograms and histograms with multiple peaks have larger CV values compared to those with a single peak. Analyzing the shape of the histogram implies that the observer introduces new interpretative elements beyond the initial observation. Therefore, even if a single peak transforms into a histogram with multiple peaks, it should be considered an increase in CV without changing the “minimal unit of recognition” or assigning any special meaning. Similarly, to preserve the “minimal unit of recognition,” whether the increase or decrease in M compared to the control is slight (minor, small, faint, mild, delicate, etc.) or major (considerable, substantial, severe, pronounced, etc.), the degree of change is not reflected; instead, only the direction of change (increase, decrease, or no change) is determined. When comparing two data points (two samples) in a single comparison, it is rare for biological phenomena to yield exactly the same values down to the decimal point unless the results are converted into integer scores, such as in a scoring system. Therefore, Type III is a rare observation. However, biological phenomena do not necessarily always fall into the same specific type. Thus, experiments must be repeated, and statistical analysis must be used to determine the differences between CV*c* and CV*e*, as well as between M*c* and M*e*, with respect to probability. This leads to cases where statistical significance is present and where it is not. For example, there may be situations in which the difference in CV is statistically significant, but the difference in M is not. When all possible combinations of statistical significance for each factor are considered, the initially proposed five patterns can be subdivided into nine distinct patterns as follows:

1. The difference in M*c* < M*e* is significant, and the difference in CV*c* > CV*e* is significant, which corresponds to Type I.
2. The difference in M*c* < M*e* is significant, and the difference between CV*c* and CV*e* is not significant, corresponding to Type I/II.
3. The difference in M*c* < M*e* is significant, and the difference in CV*c <* CV*e* is significant, corresponding to Type II.
4. The difference between M*c* and M*e* is not significant, and the difference in CV*c* > CV*e* is significant, corresponding to Type II/IV.
5. The difference in M*c* > M*e* is significant, and the difference in CV*c* < CV*e* is significant, corresponding to Type IV.
6. The difference in M*c* > M*e* is significant, and the difference between CV*c* and CV*e* is not significant, corresponding to Type IV/V.
7. The difference in M*c* > M*e* is significant, and the difference in CV*c* > CV*e* is significant, corresponding to Type V.
8. The difference between M*c* and M*e* is not significant, and the difference in CV*c* > CV*e* is significant, corresponding to Type I/V.
9. The difference between M*c* and M*e* is not significant, and the difference between CV*c* and CV*e* is not significant, corresponding to Type III. Furthermore, Type III may or may not result in a correlation between M and CV. Type III can be further divided into two cases, as follows:
10. When there is a positive correlation between M and CV, corresponding to Type II/V.
11. If there is a negative correlation between M and CV, corresponding to Type I/V.

In summary, cell behavior classification based on statistical analysis can be divided into 11 patterns. This is shown in a matrix table with M and CV as axes (Fig. 1g). These 11 patterns represent all possible cell behaviors at the minimal unit of recognition, and the method used to classify them into these 11 patterns is termed “BECC.”

### Verification of linkage between CV and cell behavior using agent-based modeling (ABM) simulations: modeling biological phenomena in ABM simulations with the presence of an observer

In this section, we demonstrate that changes in CV are universally observed phenomenon due to Attraction and Het-App. We used an ABM simulation as the method of verification. In this ABM simulation, the agent behaves according to random numbers generated at each iteration (i.e., observation). Therefore, the agent does not exhibit the same behavior in every run, which serves as a mathematical guarantee that the outcome is not artificially predetermined.

To conduct an ABM simulation, it is necessary to express cellular behavior in terms of agent behavior. Moreover, to gain consensus among observers across different spatiotemporal contexts, it is essential to use language to impart universality to what the agents represent. Therefore, this study adopted three widely recognized definitions of life. Additionally, because the presence of an observer is fundamental to this study (see Premise 2), the observer’s perspective was also incorporated. The three widely accepted definitions of life are as follows: i) being enclosed by a membrane that separates the organism from the external environment, ii) engaging in metabolism (flows of matter and energy), and iii) the ability to self-replicate.

By adding the observer’s perspective, life can be conceptually encapsulated in a single expression: “Life is that which, through cellular responses to stimuli, can be measured by an observer.” The rationale behind this expression is as follows: i) Being enclosed by a membrane can be represented as a cell. ii) Engaging in metabolism and iii) self-replication are included within responses to stimuli, because the definitions of metabolism and self-replication depend on the observer’s perspective. Thus, the phrase “measured by an observer” subsumes these processes within responses to stimuli. For example, what qualifies as a self-replica, or whether something that does not replicate during the observation period can still be called a living organism— these interpretations vary depending on the observer. Additional criteria sometimes cited in definitions of life include iv) responding to stimuli, and v) maintaining homeostasis, etc. (Gomez-Marquez 2021, Rosslenbroich 2016). These also depend on the observer’s perspective and can be unified under the linguistic expression “responses to stimuli measured by an observer.”

The definition of what constitutes a “cell” also depends on the observer. In this study, a symbol such as a circle (○), set as an agent, is considered a cell. Thus, treating ○ as a cell reflects the observer’s arbitrariness within the scope of this study. Below, we explain how and why ○ can be regarded as a “cell” in the context of being an agent.

Since an observer cannot fully recognize all aspects of a system, they cannot guarantee that an observed cell exhibits all possible reactions. Therefore, even with current research methods, it is not possible to define or establish a universal concept of the cell. In actual experimental systems, the cells under observation are arbitrarily selected by the observer.

This raises the following question: what assumptions must an observer make to ensure (or at least not deny) that an arbitrarily selected cell can exhibit all possible reactions? Upon introspection, a clear structure emerges: the observer operates under the assumption that there are more observable parameters than can be recognized at once. In other words, the observer ensures the potential for all possible cellular reactions by both acknowledging the possibility of multiple observational parameters and exercising the freedom to choose which parameters to observe. This structure, where multiple reactions exist and the observer selects a parameter at will, guarantees the possibility of all reactions for the arbitrarily selected cell.

The BECC is based on the premise that cellular reactions occur in an undefined multi-dimensional space, and from within that space, the observer freely selects an ideal one-dimensional parameter. This is structurally identical to the system that guarantees (or does not deny) that an arbitrarily selected cell can exhibit all possible reactions. The same structure applies to the circle (○) in the ABM, treating it as an agent.

The factor that causes a cell to respond is symbolized as the activator “△.” This does not represent a specific substance; rather, it encompasses physical factors and changes occurring within the cell. “△” does not represent a single molecule but represents the point of origin for factors, both internal and external, that trigger cellular responses. Its size relative to the cell and shape are irrelevant. To illustrate, this does not assume a threshold such as “three activators must encounter the cell to initiate a response.” Instead, the activator is a conceptual agent that enables the system to surpass the threshold required to initiate an appropriate response. Therefore, regardless of the specific mechanism by which a response begins, △ represents the point of initiation.

To express the behavior of an agent that reacts to stimuli, simultaneously embodying an active internal drive and passive external influence, we represented both the activator approaching the cell and the cell approaching the activator. Specifically, both the cell and activator were generated at random positions and moved randomly. Therefore, the scene can be interpreted as the cell actively moving to capture the activator or the activator moving to stimulate the cell. Thus, the system allows observation from both active and passive perspectives, depending on the observer’s perspective.

The act of “reacting” was visualized as fluctuations in a numerical value called the “response value” of a cell triggered by its encounter with an activator. Each cell was initially assigned a random response value within a certain range. While the cell retains the stimulus (or while the stimulus retains the cell), the response value increases at a random rate within a defined range. Once the retention period ends, the response value decreases at a random rate. The retention duration was randomized as well within a certain range. These changes in the response value symbolically represent measurable phenomena observed and quantified by the observer, such as phosphorylation, gene expression, image analysis, interactions, or even multicellular behavior such as predation.

Based on the explanation above, it is demonstrated that the statement “Life is that which, through cellular responses to stimuli, can be measured by an observer” can be universally applied by expressing it through agent behavior. Next, by correlating this linguistic expression with the behavior of agents, it can be represented as shown in Fig. 2a.

Finally, the modeling was based on finiteness. Infinity is a significant concept in philosophy, mathematics, and physics, and assuming infinity can be a meaningful approach. However, while it is possible to treat concepts such as “infinitely existing cells” or “infinite reactions” in theory, we have not encountered any experimental observations in biological phenomena where one could realistically collect an infinite number of cells or continuously measure infinite reactions.

If one assumes that life phenomena can persist indefinitely, then an end state like the Big Freeze must be avoided. However, current cosmological models suggest that the universe is headed towards an open, ever-expanding state that will ultimately result in a Big Freeze. Therefore, the assumption that life phenomena can continue forever is not supported by current scientific understanding of the universe’s ultimate fate.

Therefore, in this study, we adopted experimental conditions under the assumption of finite resources, and we did not presume that the number of cells or response values were infinite. Accordingly, the biological responses represented as agents in this study are finite, and, these responses are defined to have both a beginning and an end.

### Verification of linkage between CV and cell behavior using ABM simulations: observation procedures for BECC with ABM simulations

“Being measured by the observer” refers to assessing the reactions occurring within the target cell population and the magnitude of those reactions, as represented by changes in the M and CV in an ideal one-dimensional histogram. The reason for using M and CV is that they constitute the minimal unit of recognition, as explained in Fig. 1.

The M is calculated as the sum of the individual response values (R1…Rn) for each of the n cells in the population, divided by n:

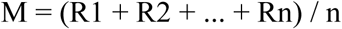

The CV is derived from the histogram of these individual response values. To minimize the influence of sample size and avoid assuming a normal distribution, we used a robust CV calculated using the median and percentiles. This method follows formulas similar to those used in flow cytometry, a device commonly employed to measure individual cells. The formula is as follows.

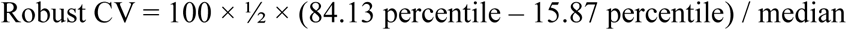

Several methods exist for calculating percentiles. In this simulation, we adopted an exclusive method excluding 0 and 1 when determining the total rank range. As described later, the number of cells forming the population in this simulation was set to approximately 1,000, providing a sufficiently large sample size for the percentile calculation. Thus, the differences in the calculated percentile values due to the choice of method were negligible.

The observer is the entity that defines an ideal one-dimensional parameter and an observation range, and then analyzes the resulting data. In the ABM simulation, the response value serves as the value of an ideal one-dimensional parameter. Therefore, the act of setting the remaining simulation conditions and initiating the simulation to obtain CV and M values constitutes the definition of the observation, and then the entity that analyzes the resulting numerical data is regarded as the observer.

In this study, ABM simulations were conducted using the NetLogo programming environment. NetLogo is widely adopted and user-friendly, even for those who are not highly proficient in programming, which increases the likelihood of verification by observers with different spatiotemporal perspectives. For these reasons, NetLogo was selected as the programming platform for this study. The NetLogo program file (Model of Cell Population ver. 5.0.2) used in this study and its program code are provided as Supplementary Information. The representative interface tab is shown in Fig. 2b.

### Verification of linkage between CV and cell behavior using ABM simulations: initial response value and CV

Observation begins with a given result, which is the same as deciding on an arbitrary cell by the observer. That is, the initially assigned conditions of “attraction” and “Het-App” determine the initial response value, from which the simulation begins. In this ABM simulation, the CV of the initial response value is defined by the settings of the sliders “basal-response” and “basal-noise”, and it is determined when the “set**”** button is pressed. This process corresponds to the deliberate selection of a cell population for observation.

As shown in Fig. 3a, when the initial response value is set high, CV decreases. Conversely, even when the response value remains constant, increasing noise results in a higher CV. However, when the response value is high, an increase in noise has a limited effect on CV. When the initial state of “attraction” is strong, the effect of attraction becomes less observable. Conversely, when the initial “attraction” is weak, the effect of “heterogeneous-cell presence (Het-App)” becomes less observable.

**Fig. 3.**
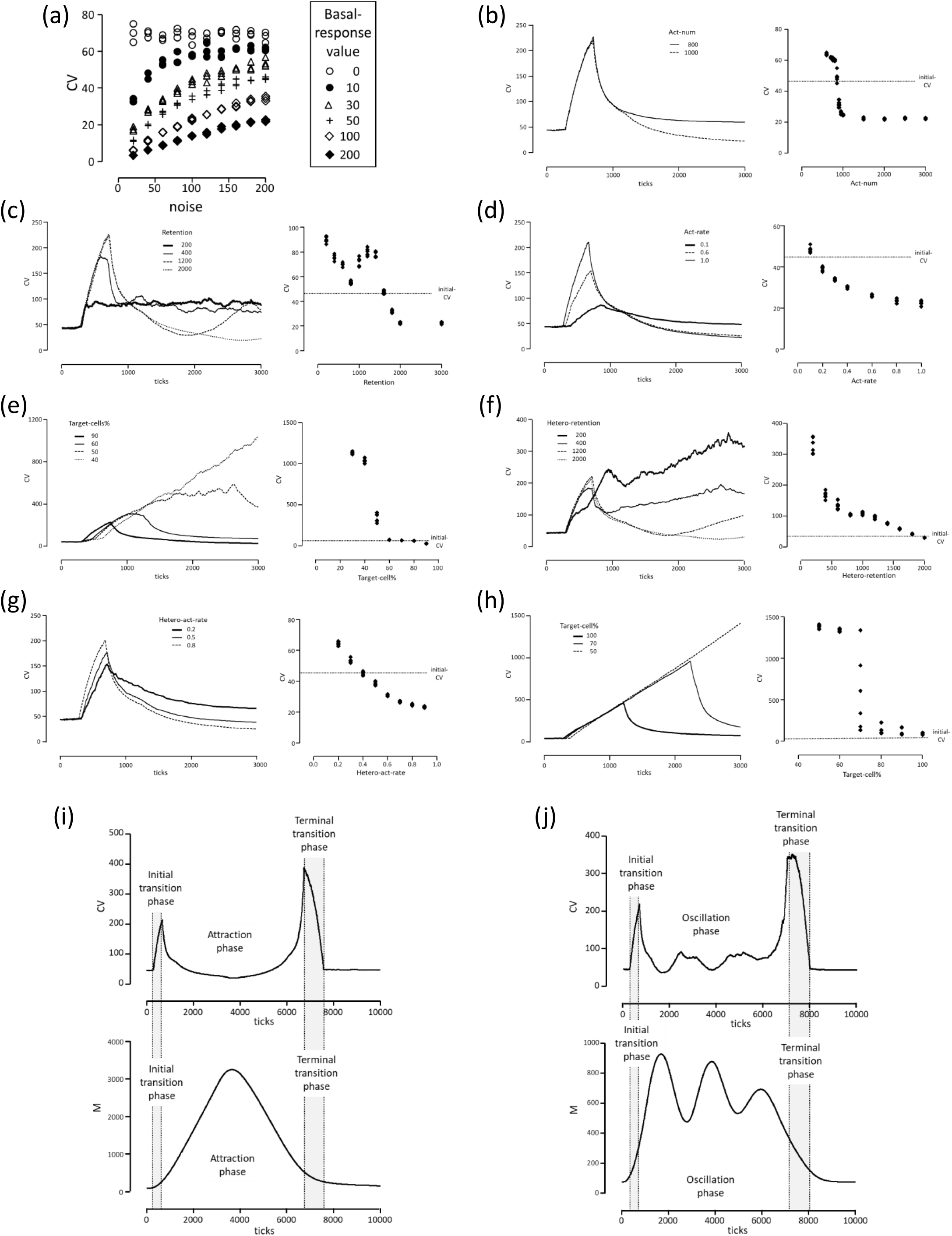
Verification of BECC using ABM simulation. (a) Initial response values and CV changes. Each Basal-Response value (0, 10, 30, 50, 100, and 200) that is initially (before the start of the simulation) set for a cell is indicated by the difference between each symbol (○, ●, △, +, ◇, and ◆). The noises on the horizontal axis indicate the range of initial response values randomly given to individual cells. The results are shown for each of the three times under the conditions of each initial reaction value setting. (b) Relationship between frequency of response (Act-num) and CV. A parameter sweep simulation was conducted under the following conditions: Total-cell-num: 1,000; Target-cell-%: 100; Finish-ticks: 3,000; Act-num: 600, 700, 750, 800, 850, 900, 950, 1,000, 1,500, 2,000, 2,500, and 3,000; basal-response: 30; basal-noise: 100; act-rate: 1.0; act-rate-noise: 0.5; deact-rate: 1.0; deact-rate-noise: 0.5; retention: 3,000; retention-noise: 50. The results of three independent runs of this parameter sweep are shown.(c) Relationship between strength of response (Act-rate) and CV. The parameter sweep simulation was conducted under the following conditions: Total-cell-num: 1,000; Target-cell-%: 100; Finish-ticks: 3,000; Act-num: 3,000; basal-response: 30; basal-noise: 100; act-rate: 0.1, 0.2, 0.4, 0.6, 0.8, 1.0; act-rate-noise: 0.5; deact-rate: 1.0; deact-rate-noise: 0.5; retention: 3,000; retention-noise: 50. (d) Relationship between response-retention of Het-App (Heter-retention) and CV. The parameter sweep simulation was conducted under the following conditions: Total-cell-num: 1,000; Target-cell-%: 50; Finish-ticks: 3,000; Act-num: 3,000; basal-response: 30; basal-noise: 100; act-rate: 1.0; act-rate-noise: 0.5; deact-rate: 1.0; deact-rate-noise: 0.5; retention: 3,000; retention-noise: 50; Hetero-cells-response-to-act: ON (true); hetero-act-rate: 1.0; hetero-act-rate-noise: 0.5; hetero-deact-rate: 1.0; hetero-deact-rate-noise: 0.5; hetero-retention: 200, 400, 600, 800, 1000, 1,200, 1,400, 1,600, 1,800, and 2,000; hetero-retention-noise: 50. (e) Relationship between proportion of Het-App (Target-cell%) and CV. The parameter sweep simulation was conducted under the following conditions: Total-cell-num: 1,000; Target-cell-%: 90, 80, 70, 60, 50, 40; Finish-ticks: 3,000; Act-num: 3,000; basal-response: 30; basal-noise: 100; act-rate: 1.0; act-rate-noise: 0.5; deact-rate: 1.0; deact-rate-noise: 0.5; retention: 3,000; retention-noise: 50; Hetero-cells-response-to-act: ON (true); hetero-act-rate: 0.1; hetero-act-rate-noise: 0.5; hetero-deact-rate: 0.1; hetero-deact-rate-noise: 0.5; hetero-retention: 200; hetero-retention-noise: 50. (f) Relationship between response-retention of Het-App (Heter-retention) and CV. The parameter sweep simulation was conducted under the following conditions: Total-cell-num: 1,000; Target-cell-%: 50; Finish-ticks: 3,000; Act-num: 3,000; basal-response: 30; basal-noise: 100; act-rate: 1.0; act-rate-noise: 0.5; deact-rate: 1.0; deact-rate-noise: 0.5; retention: 3,000; retention-noise: 50; Hetero-cells-response-to-act: ON (true); hetero-act-rate: 1.0; hetero-act-rate-noise: 0.5; hetero-deact-rate: 1.0; hetero-deact-rate-noise: 0.5; hetero-retention: 200, 400, 600, 800, 1000, 1,200, 1,400, 1,600, 1,800, and 2,000; hetero-retention-noise: 50. (g) Relationship between response-retention of Het-App (Heter-act-rate) and CV. The parameter sweep simulation was conducted under the following conditions: Total-cell-num: 1,000; Target-cell-%: 50; Finish-ticks: 3,000; Act-num: 3,000; basal-response: 30; basal-noise: 100; act-rate: 1.0; act-rate-noise: 0.5; deact-rate: 1.0; deact-rate-noise: 0.5; retention, 3000; retention-noise: 50; Hetero-cells-response-to-act: ON (true); hetero-act-rate: 0.2, 0.3, 0.4, 0.5, 0.6, 0.8, and 0.9; hetero-act-rate-noise: 0.5; hetero-deact-rate: 0.1; hetero-deact-rate-noise: 0.5; hetero-retention: 3,000; hetero-retention-noise: 50. (h) Relationship between non-response of heterogenous cells and CV. The parameter sweep simulation was conducted under the following conditions: Total-cell-num: 1,000; Target-cell-%: 50 60 70 80 90 100; Finish-ticks: 3,000; Act-num: 500; basal-response: 30; basal-noise: 100; act-rate: 1.0; act-rate-noise: 0.5; deact-rate: 1.0; deact-rate-noise: 0.5; retention: 3,000; retention-noise: 50; Hetero-cells-response-to-act: OFF (false). (b – h) The graphs on the left-side presents representative examples of each parameter-sweep condition as shown in each line (thick line, solid line, thick dashed line, or solid dashed line). The graphs on the right-side shows a summary of the results at the observation endpoint of 3,000 ticks, and the dashed line indicates the initial CV caused by the reaction values set as initial conditions. The results of three independent runs of this parameter sweep are shown. (i) Relationship between time lapses and CV with attraction phase. The simulation parameters were conducted under the following: Total-cell-num: 1,000; Target-cell-%: 100; Finish-ticks: 10,000; Act-num: 1,000; basal-response: 30; basal-noise: 100; act-rate: 1.0; act-rate-noise: 0.5; deact-rate: 1.0; deact-rate-noise: 0.5; retention: 3,000; retention-noise: 50. The range of ticks from when the CV starts to increase to when it begins to decrease was defined as the *initial transition phase* (300–599 ticks). Additionally, the range of ticks from the point at which M and CV return to their pre-reaction values, going backward to the tick at which the CV reaches its maximum, was defined as the *terminal transition phase* (7,000–7,600 ticks). The interval between the initial and terminal transition phases (600–6,999 ticks) was defined as the *attraction phase*. The upper graphs show the results of CV, and the lower graphs are the mean (average) of response value (M). Each initial transition phase and terminal transition phase are presented as dashed lines and light gray areas. These results are representative of three independent runs. (j) Relationship between time lapses and CV with oscillation phase. The simulation parameters were conducted under the following: Total-cell-num: 1,000; Target-cell-%: 100; Finish-ticks: 10,000; Act-num: 3,000; basal-response: 30; basal-noise: 100; act-rate: 1.0; act-rate-noise: 0.5; deact-rate: 1.0; deact-rate-noise: 0.5; retention: 1,000; retention-noise: 50. The range of ticks from when the CV starts to increase to when it begins to decrease was defined as the *initial transition phase* (300–700 ticks). Furthermore, the range of ticks from the point at which M and CV return to their pre-reaction values, going backward to the tick at which the CV reaches its maximum, was defined as the *terminal transition phase* (6,800–8,000 ticks). The interval between the initial and terminal transition phases (701–6,799 ticks) was defined as the *oscillation phase*. The upper graphs show the results of CV, and the lower graphs are the mean (average) of response value (M). Each initial transition phase and terminal transition phase are presented as dashed lines and light gray areas. These results are representative of three independent runs.

Since the purpose of this paper is to simulate and observe both “attraction” and “Het-App” as results of cellular stimulation, the simulations that follow were conducted under the condition that both effects can be clearly observed. The initial settings were basal-response: 30 and basal-noise: 100.

### Verification of linkage between CV and cell behavior using ABM simulations: simulation of “Attraction”

In the thought experiment, it was assumed that a decrease in CV results from “Attraction,” which is defined as a set of effects that cause a cell to remain in a particular state. The effects that constitute attraction are classified into three integrated categories: i) frequency of response, ii) duration of response, and iii) strength of response. In this ABM simulation, these are respectively represented by i) Act-num, ii) Retention, and iii) Act-rate. When the values of these parameters are higher in the experimental group than in the control group, it is considered that attraction has occurred. This simulation examined whether CV decreases under such conditions.

Initial response value and CV were decided as described above (Fig. 3a). To clarify the effect of attraction on decreasing CV, the frequency of response (Act-num) was examined. The number of activators appearing (Act-num) varied from 600 to 3,000 to compare changes in CV. All other conditions were kept constant. The simulation conditions (Act-num parameter sweep) are described in Fig. 3b. Representative examples of temporal changes are shown for Act-num, values of 800 and 1,000 (Fig. 3b, left). A notable pattern was a transient increase in CV observed before 1,000 ticks, where each tick corresponds to one execution of behavior by every agent according to the program code (i.e., an execution unit). This early-phase increase will be discussed later in the time-course section. Focusing on the observation point (3,000 ticks), it was found that CV was lower at Act-num 1,000 compared to Act-num 800. Further analysis showed that CV began to decrease when Act-num exceeded the total number of cells (1,000). This implied that when the frequency of reactions exceeded the total number of cells, reactions occurred in all cells and CV decreased (Fig. 3b, right). Thus, it was confirmed that CV decreases depending on the frequency of response.

To clarify the decrease in CV due to attraction, the retention of the response was examined. The response retention time (Retention) varied from 200 to 3,000 ticks and the variation of CV was compared. All other conditions were kept constant. The simulation conditions (Retention parameter sweep) are described in Fig. 3c. The observation endpoint was set at 3,000 ticks, where the response persists (i.e., the Act-num does not diminish). Representative examples of temporal changes are shown for retention durations of 200, 400, 1,200, and 2,000 ticks. A notable trend in the temporal dynamics was a transient increase in CV observed when the retention exceeded 1,000 ticks, particularly within the first 1,000 ticks. In contrast, for shorter retentions of 200 and 400, the increase in CV reached a steady state. This initial CV increase in the early phase of the temporal changes will be described later in the time-course section. A key feature of sustained responses was that in cases where the initial response was maintained until the observation point (retention 2,000 and 3,000 ticks), the CV decreased compared to the initial CV (Fig. 3c). On the other hand, when the reaction repeatedly terminated and restarted (retention ≤ 1,000 ticks), the CV increased compared to the initial value. These results confirm that when cellular responses to stimuli are sustained, the synchrony within the cell population increases, leading to a reduction in CV.

To clarify the decrease in CV due to attraction, the intensity of the response was examined. The strength of the response (Act-rate) varied from 0.1 to 1 to compare variations in CV. All other conditions were kept constant. The simulation conditions (Act-rate parameter sweep) are described in Fig. 3d. Representative examples of transitional changes are shown for Act-rate values of 0.1, 0.6, and 1.0. A notable feature of the transitional change was a transient increase in the CV observed before 1,000 ticks. This initial increase in CV during the transitional phase will be described in the time-course section. A characteristic feature of the effect of response strength was that at the observation point, there was no discontinuity, and the CV decreased in an Act-rate-dependent manner compared to the initial CV (Fig. 3d). From this, it was confirmed that when the response to a stimulus is strong at the observation point, the CV decreases. Based on these results and those presented in Fig. 3b-3d it was verified that the CV decreases with “Attraction.”

### Verification of linkage between CV and cell behavior using ABM simulations: simulation of “Het-App”

In the thought experiment, it was assumed that the increase in CV results from the appearance of heterogeneous cells (Het-App). Here, it was verified whether this increase in CV was indeed caused by Het-App, which refers to the appearance of cells that exhibit responses different from those of the main cell population. The factors constituting the impact of this emergence were classified into three integrated categories: i) the proportion of heterogeneous cells, ii) the difference in response of heterogeneous cells, and iii) non-responsiveness of heterogeneous cells. In the case of non-responsiveness, the cells do not encounter the stimulus (the stimulus is not consumed). In this ABM simulation, the corresponding parameters are: i) Target-cell% for the proportion, ii) Hetero-act-rate and Hetero-retention of heterogenous cells (Hetero-cells) for the difference in response, and iii) the OFF (false) state of the switch for Hetero-cells-response-to-act for non-responsiveness. When the values of these parameters were higher in the experimental group than in the control group, heterogeneous cells were considered to have emerged.

To verify that Het-App led to an increase in CV, the proportion of Het-App was examined. For Target-cells, the previously discussed conditions were used in which CV decreased due to Activator (retention: 3,000; act-rate: 1.0). For Hetero-cells, conditions were set in which CV did not decrease in response to the stimulus (retention: 200; act-rate: 0.1). All other conditions were kept constant. The simulation conditions (Target-cell% parameter sweep) are described in Fig. 3e. The activator was configured to interact with both Target-cells and Hetero-cells, and the Act-num was set to 3,000 to ensure sustained responses. A representative example of temporal changes is shown in Fig. 3e, illustrating varying proportions of heterogeneous cells with Target-cell% at 90, 60, 50, and 40%. A notable pattern in these temporal changes was that when the proportion falls below 50%, the transient increase in CV was no longer followed by a decrease, indicating a persistent loss of synchrony. Another distinctive observation related to the proportion of Het-App was that at the observation endpoint 3,000 ticks, the CV did not fall below its initial value once the Target-cell% dropped below 90%, and it increased markedly when the percentage fell below 50%. These findings confirm that CV increases depending on the proportion of Het-App.

To verify that differences in the responses of heterogeneous cells led to an increase in CV, variations in response duration were examined with heterogeneous cells present at a fixed proportion. The proportion of Target-cells was fixed at 50%, using the previously discussed condition in which CV decreased due to activator (retention: 3,000 ticks). For Hetero-cells, the duration of response was varied, with retention values ranging from 200 to 2,000 ticks. All other conditions were kept constant. The simulation conditions (Hetero-retention parameter sweep) are described in Fig. 3f. The observation endpoint was set at 3,000 ticks, where the response persists (i.e., the Act-num does not diminish). The activator was set at Act-num 3,000, a condition that met both Target-cells and Hetero-cells, sustaining the response. A representative example of temporal changes is shown in Fig. 3f, where the proportion of heterogeneous cells is fixed at 50%, and Hetero-retention is set at 200, 400, 1,200, and 2,000 ticks. The hetero-retention value decreases, indicating a divergence in the properties of the hetero-cell and target-cell (with a retention value of 3000), which leads to increased heterogeneity. A notable trend in these temporal changes was that when Hetero-retention is short (200 or 400 ticks), the decrease in CV was not maintained and instead increased. Another key observation regarding response differences was that at the point of observation, when the hetero-retention of Hetero-cells falls below 400 ticks, CV increases significantly. This suggests that the pronounced increase in CV resulted from the divergence between the periodic fluctuations of Hetero-cells and the sustained responses of Target-cells. These results confirm that when the maintenance of response to stimuli in Hetero-cells differed from that in Target-cells, CV increased at the point of observation.

To verify that differences in response strength among heterogeneous cells led to an increase in CV, variations in response strength were examined with heterogeneous cells present at a fixed proportion. The proportion of Target-cells was fixed at 50%, using the previously discussed condition in which CV decreased due to activator (act-rate: 1.0). For Hetero-cells, the response strength was set to values different from that of Target-cells (Hetero-act-rate: 0.2 to 0.9). All other conditions were kept constant. The simulation conditions (Hetero-act-rate parameter sweep) are described in Fig. 3g. The activator was also configured to interact with both Target-cells and Hetero-cells, and the Act-num was set to 3,000 to ensure sustained responses. A representative example of temporal changes is shown in Fig. 3g, with Hetero-act-rate values set to 0.2, 0.5, and 0.8. The Hetero-act-rate value decreases, indicating a divergence in the properties of the hetero-cell and target-cell (with an act-rate value of 1.0), which leads to increased heterogeneity. A notable trend in these temporal changes was that when the Hetero-act-rate diverges significantly from the Target-cell act-rate of 1.0, specifically at 0.2, the transient increase in CV was not followed by a decrease below the initial CV level. A key observation regarding differences in response strength was that at the observation point, divergence between the Hetero-act-rate and the Target-cell act-rate of 1.0 resulted in an increase in CV (Fig. 3g). These results confirm that when the response strength of Hetero-cells to stimuli differs from that of Target-cells, CV increases at the point of observation.

Next, the conditions were examined in which Hetero-cells did not respond at a given proportion. One could interpret this as equivalent to a “Hetero-act-rate” of 0 in terms of reaction strength. However, a key difference was that when Hetero-cells encounter activators, the activators were not consumed. This suggests the possibility that the ratio of Target-cells to activators might increase. Therefore, the condition was fixed at Act-num 500, and cases were investigated in which the Target-cell percentage ranged from 50% to 100%. All other conditions were kept constant. The simulation conditions (Target-cell-% parameter sweep on Hetero-cells-response-to-act, OFF) are described in Fig. 3h. Representative examples of temporal changes are shown for Target-cell% values of 50%, 70%, and 100% (Fig. 3h). A notable feature of these transitional changes is that, unlike the 100% Target-cell case, the coefficient of variation (CV) continues to increase in the 50% case (Fig. 3h). From this, it is confirmed that the presence of non-reactive Hetero-cells led to an increase in CV. These results, in conjunction with the data shown in Fig. 3e-3h, confirm that “Het-App” leads to an increase in CV.

### Verification of linkage between CV and cell behavior using ABM simulations: simulation of “time”

As previously mentioned, we examined whether CV fluctuates during the transition process as a result of time-dependent state changes. It has already been shown in the simulation results concerning attraction and Het-App that a transient increase in CV occurs during the initial phase of the reaction, as illustrated by representative examples of transitional changes. Therefore, we confirmed that fluctuations in CV also occur during the late stages of the reaction as part of the overall reaction process. Here, to explicitly indicate the start and end of the entire reaction process, we also show the temporal changes in the mean value of the reaction, M. Specifically, under the simulation conditions for attraction, the observation period was extended up to 10,000 ticks. As a representative example, we extended the observation period up to 10,000 ticks under the condition of sufficient attraction (retention 3,000; Fig. 3i). As a result, as shown in Fig. 3i, a transient increase in CV occurred in the early phase of the reaction (between 300 and 599 ticks), when the reaction value began to rise. Subsequently, CV decreased in parallel with the increase in the reaction value. In the late phase of the reaction, when the reaction value returned to its initial level (between 7,000 and 7,600 ticks), another transient increase in CV was observed. Furthermore, under the condition of retention 1,000 (Fig. 3j), where the reaction becomes oscillatory, the observation period was also extended to 10,000 ticks. As shown in Fig. 3j, a similar increase in CV was observed during both the initial phase of the reaction and its concluding phase. From these results, it was confirmed that CV increases during the transitional phases, both the early and late stages, when reactive and non-reactive cells coexist within the cell population. Through the above simulations (Fig. 3i and 3j), it was demonstrated that Het-App and attraction, occurring temporally, lead to both increases and decreases in CV.

### Proof of universality of BECC using ABM simulation results: comparison at observation points between two different conditions

We examined the case where cells were stimulated and compared at a certain observation point (see Confirmation 5). The stimulation was conducted under the conditions described in Fig. 3b, and both cases, where the CV increased and where it decreased, were investigated (Fig. 4a). When CV increased at the observation point: In the observation comparing Act-num 0 (Observation Point A) and Act-num 800 (Observation Point B), with Observation Point A as the control group (lower subscript *c*) and Observation Point B as the experimental group (lower subscript *e*), both M and CV increased. Therefore, this was classified as Type II (M↑, CV↑) (Fig. 4b; A*c*-B*e*).

**Fig. 4.**
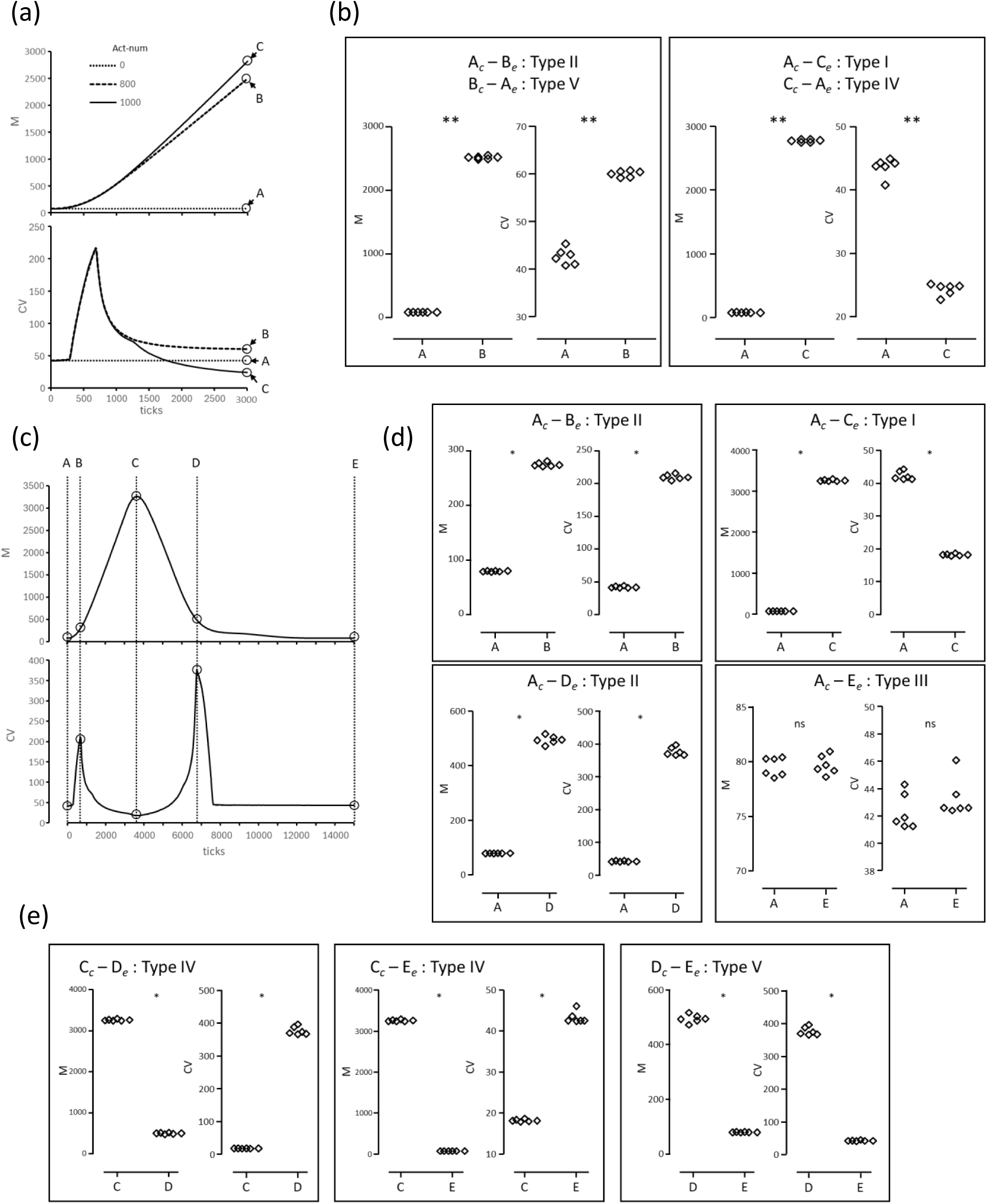
Verification of BECC using ABM simulation. (a) Representative example of observation point comparison in cellular responses. The ABM simulation was conducted under the same conditions as in Fig. 3b. As representative examples, Act-num values of 0 (thin dashed line), 800 (thick dashed line), and 1,000 (thin solid line) are shown. The upper graph displays the results of the response value M, and the lower graph shows the results of CV. The points indicated by circles and arrows represent observation points A, B, and C at 3,000 ticks. (b) BECC based on observation point comparison in cellular responses. Simulations were conducted under the same conditions as in Fig. 1c. Observation points A, B, and C shown in Fig. 4a were respectively compared as control groups (subscript-italic *c*) and experimental groups (subscript-italic *e*). Each diamond-shaped symbol represents M (left panel) and CV (right panel) obtained from six independently conducted simulations. The horizontal axis indicates the observation points. Statistical analysis was performed using Mann-Whitney U test. ns, non-significant; *, *p* < 0.05; **, *p* < 0.01. (c) Representative example of time-course comparison in cellular responses. Under the same conditions as in Fig. 3i, observations were made up to 10,000 ticks. The upper graph shows the results of the response value M, and the lower graph shows the results of CV. Characteristic fluctuations in CV are indicated by points A – E, each shown as intersections of circles and dashed lines. (d) BECC based on time-course comparison in cellular responses: Control group is Observation Point A. Simulations were conducted under the same conditions as in Fig. 3i, up to 10,000 ticks. Observation point A shown in Fig. 4c was used as the control group (subscript-italic *c*), and B – E were compared as experimental groups (subscript-italic *e*). (e) BECC based on time-course comparison in cellular responses: Control groups are Observation Points C or D. Simulations were conducted under the same conditions as in Fig. 3i, up to 10,000 ticks. Observation points C or D shown in Fig. 4c were used as control groups (subscript-italic *c*), and D and E were compared as experimental groups (subscript-italic *e*). (B, D, and E) Each diamond-shaped symbol represents the M (left panel) and CV (right panel) obtained from six independently conducted simulations. The horizontal axis indicates the observation points. Statistical analysis was performed using the Mann-Whitney U test. ns, non-significant; *, *p* < 0.05; **, *p* < 0.01.

When the roles were reversed, with Observation Point B as the control group (lower subscript *c*) and Observation Point A as the experimental group (lower subscript *e*), such as in a scenario where an inhibitor was added, both M and CV decreased. This was classified as Type V (M↓, CV↓) (Fig. 4b; B*c*-A*e*). Conversely, when CV decreased at the observation point: In the observation comparing Act-num 0 (Observation Point A) and Act-num 1,000 (Observation Point C), with Observation Point A as the control group (lower subscript *c*), the experimental group (lower subscript *e*) at Observation Point C showed an increase in M and a decrease in CV. This was classified as Type I (M↑, CV↓) (Fig. 4b; A*c*–C*e*). Furthermore, when Observation Point C served as the control group and Observation Point A as the experimental group, M decreased and the CV increased. This was classified as Type IV (M↓, CV↑) (Fig. 4b; C*c*–A*e*).

### Proof of universality of BECC using ABM simulation results: comparison of different time points under the same condition

Cells were stimulated under conditions that induced a decrease in CV at 3,000 ticks (conditions from Fig. 3b), and observations were continued up to 10,000 ticks (Fig. 4c). Based on the results, representative observation points were selected at which CV showed characteristic fluctuations, namely, Observation Points A, B, C, D, and E, and comparisons were made among these points. Since this process represents a unidirectional and irreversible event, the Observation Points shown left-side in the figures. serve as the control groups (see Confirmation 5). After stimulation, the CV of the cell population exhibited transient changes: an increase (Observation Point B), a decrease (Observation Point C), another increase (Observation Point D), and a final decrease (Observation Point E). In parallel, M increased to a peak at Observation Point C, then gradually decreased toward Observation Point E (Fig. 4c). This sequence represents the initiation and termination of a cellular response. When Point A was set as the control group, the progression followed a sequence of Type II → I → II → III (Fig. 4d; A*c*-B*e*, A*c*-C*e*, A*c*-D*e*, A*c*-E*e*). Furthermore, when the peak point of M (Point C) was used as the control group or when Observation Point D (where M begins to decrease) was used as the control group and Observation Point E (the endpoint of the response) was used as the experimental group, the sequence became Type IV → V (Fig. 4e; C*c*-D*e*, C*c*-E*e*, D*c*-E*e*).

The transitions of each BECC type can be described as follows:

A*c*-B*e*: Type II (M↑, CV↑) indicates an increase in heterogeneous cells due to the response of the cell population when the observation point of the control group is set before the onset of the response.

A*c*-C*e*: Type I (M↑, CV↓) suggests attraction due to the peak phase of the response when the observation point of the control group is set before the onset of the response.

C*c*-D*e*: Type IV (M↓, CV↑) shows an increase in heterogeneous cells due to a decrease in response when the observation point of the control group is set at the peak phase of the response.

D*c*-E*e*: Type V (M↓, CV↓) suggests attraction due to the termination of the response when the observation point of the control group is set just before the end of the response.

A*c*-E*e*: Type III (M→, CV→) indicates there is no difference in the cell population after the end of the response when the observation point of the control group is set before the onset of the response.

### Proof of universality of BECC using ABM simulation results: Type III – initial/terminal transition phase and attraction/oscillation phases

Even in the presence of observer-induced experimental error (see Confirmation 7), if both M and CV show statistically significant differences, cell behavior can still be classified as Type I, II, IV, or V, as described earlier. If neither shows a significant difference, the behavior is classified as Type III. However, observer-induced experimental error can sometimes lead to a Type III classification. A Type III classification (M→, CV→) may indicate that the observed region corresponds to a specific behavioral state of the cell population where the cumulative response values over a certain range of ticks exhibit correlation.

One specific example of such a special condition can be found in the simulation results shown in Fig. 3i and 3j, particularly during the initial transition Phase and the terminal transition phase. During these periods, both M and CV fluctuate. When data from these fluctuating phases are aggregated, the resulting values exhibit large variability. Consequently, it becomes difficult for significant differences to be detected in the cumulative results of such an observation region. If no significant difference is detected, one can examine the correlation between M and CV by combining the results of the control and experimental groups. This allows detection of the positive correlation observed during the initial or terminal transition phases (Fig. 5a). This is because, as the response M changes, heterogeneous cells emerge, leading to an increase in CV. This phase can be described as a positive correlation (M ↔ CV, *r* > 0), corresponding to Type II/V. While it may not be possible to distinguish between the initial and terminal transition phases, it becomes evident that the observed cells are in a transitional phase.

**Fig. 5.**
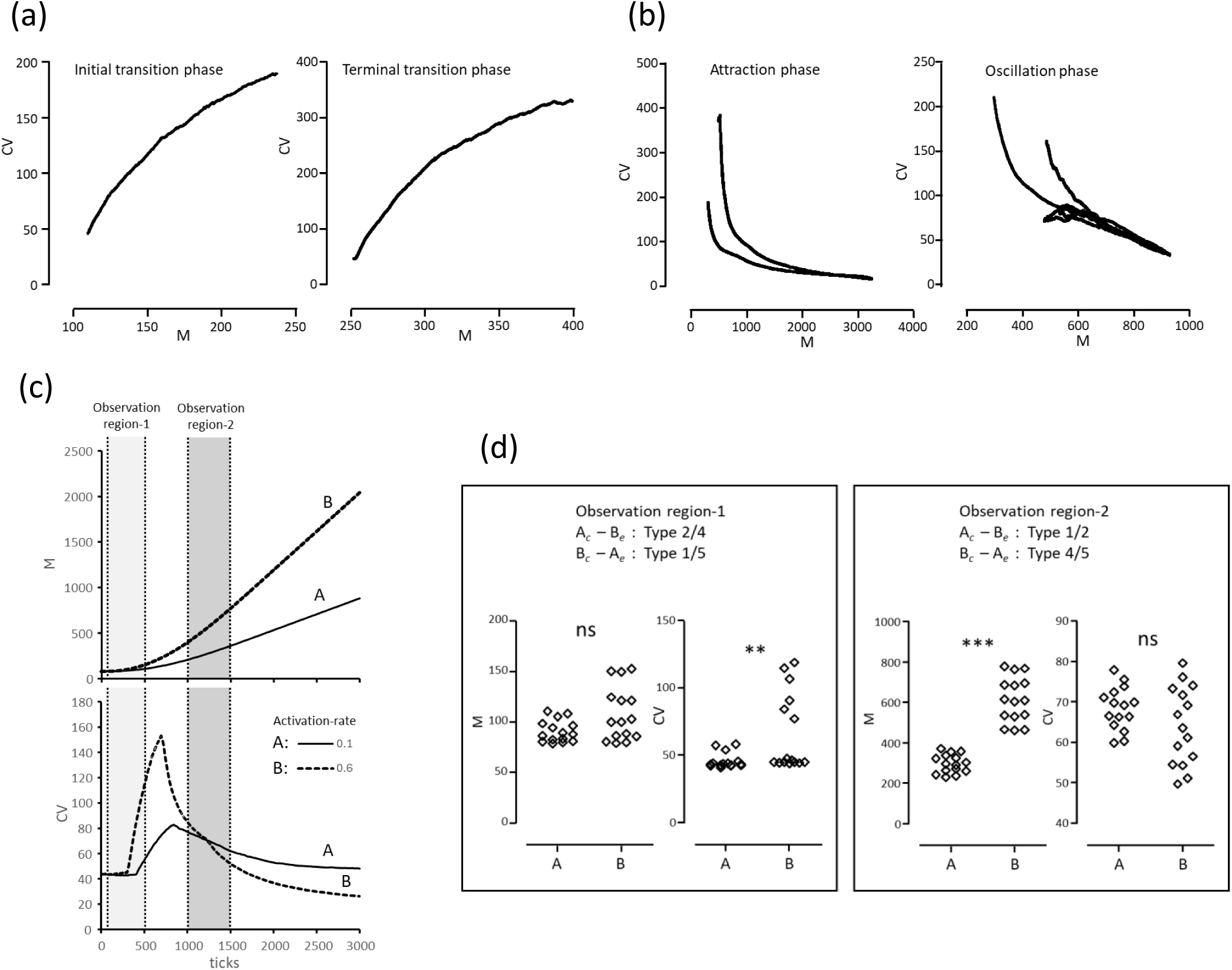
Validation of BECC using ABM simulations in cases where inter-experimental error exceeds the biological fluctuation of the phenomenon. (a) A positive correlation between M and CV when the observation range consists of results from the transition phase. In the simulation results shown in Fig. 3i, the left panel illustrates the correlation between M and CV during the initial transition phase, while the right panel illustrates the correlation during the terminal transition phase. In both cases, the correlation coefficient was *r* = 1.0, *p* < 0.0001. (b) A negative correlation between M and CV when the observation range consists of results from the attraction or oscillation phase. The left panel shows the correlation between M and CV during the attraction phase in the simulation results from Fig. 3i (*r* <-0.98, *p* < 0.0001). The right panel shows the correlation during the oscillation phase in the simulation results from Fig. 3j (*r* <-0.86, *p* < 0.0001). (A and B) The correlation coefficient *r* and *p*-value were calculated using Spearman rank test. (c) A representative example where the observation range results in no significant difference in either M or CV. The simulation shown is part of the results presented in Fig. 3d. It displays the outcomes for Act-rate 0.1 (A, thin solid line) and Act-rate 0.6 (B, thick dashed line). Observation region-1 (light gray) was arbitrarily set to 0–500 ticks, and Observation region-2 (dark gray) to 1,000–1,500 ticks. (d) BECC based on the results from Observation region-1 and-2 as shown in (c). Three independent simulations were conducted under the conditions described in Fig. 3d. From the results that fell within the range of Observation region-1, data at 100, 200, 300, 400, and 500 ticks were selected, and M and CV were summarized in separate graphs (left panel). Similarly, from the results within the range of Observation region-2, data at 1,100, 1,200, 1,300, 1,400, and 1,500 ticks were selected, and M and CV were summarized in separate graphs (right panel). On the horizontal axis, labels A and B indicate different Act-rate conditions: 0.1 and 0.6, respectively. Statistical analysis was performed using Mann-Whitney U test. ns, non-significant; **, *p* < 0.01; ***, *p* < 0.001.

Another special condition can be seen in the simulation results shown in Fig. 3i and 3j during the attraction phase and oscillation phase. In these periods, the states of attraction and the appearance of heterogeneous cells coexist, causing variations in both M and CV. When the system is in the attraction phase or oscillation phase, attraction occurs as part of the response, leading to a decrease in CV. Consequently, as shown in Fig. 5b, the relationship between M and CV shows a negative correlation (M↔ CV, *r* < 0). This can be detected as Type I/IV.

### Proof of universality of BECC using ABM simulation results: comparison in observation ranges where M or CV is unclear

In typical experimental systems, it is difficult to distinguish whether inter-experiment error originates from biological fluctuations or from the observer’s influence. This issue cannot be resolved simply by increasing statistical precision; once the results are observed, the errors are inseparably mixed. In contrast, in ABM simulations of biological phenomena, these two types of errors can be separated and handled independently. When this ABM simulation is used to represent the observer’s measurement error, the observation results in an arbitrary range of ticks. (see Confirmation 7).

As a specific example in which neither M nor CV shows a significant difference, we cite the simulation results conducted under the conditions shown in Fig. 3d. In the observation region (Observation Region-1) shown in Fig. 5c, when comparing act-rate 0.1 (Condition A) and act-rate 0.6 (Condition B), there is no significant difference in M. At this point, it is possible that the increase or decrease in heterogeneous cells has already occurred. As shown in Observation Region-1, Fig. 5d; A–B, Type II/IV (M→, CV↑) indicates that the response M fluctuates beyond the observation region, and heterogeneous cells may appear. Conversely, when the control groups are reversed for comparison (Observation Region-1, Fig. 5d; B–A), it is classified as Type I/V (M→, CV↓), where the response M also fluctuates beyond the observation region, and attraction may occur. This represents a peripheral state of the main response within the cell population.

As in the previous example, we cite the simulation results conducted under the conditions shown in Fig. 3d. In the observation region (Observation Region-2) shown in Fig. 5c, when comparing act-rate 0.1 (Condition A) and act-rate 0.6 (Condition B), fluctuations in CV are large, and no significant differences are detected compared to the control group. At this point, an increase or decrease in M may occur. As shown in Observation Region-2, Fig. 5d; A– B, Type I/II (M↑, CV→) indicates that CV decreases beyond the observation region, and attraction may arise. Conversely, when the control groups are reversed for comparison (Observation Region-2, Fig. 5d; B–A), it is classified as Type IV/V (M↓, CV→), where CV increases beyond the observation region, and heterogeneous cells may appear. This represents a peripheral state of either attraction or the emergence of heterogeneous cells.

From the analysis of cumulative results from the observation region, a significant difference in only one of M or CV represents a peripheral state where a biological phenomenon may be occurring. However, this peripheral state alone doesn’t allow for the prediction of changes. Since prediction relies on temporal observation, a change in conditions might exceed the minimal unit of recognition, causing the description to shift from recognition to understanding.

In some cases, significant differences obtained by comparing ideal responses (see Confirmation 4), rather than using BECC, might reflect peripheral states of change in the cell population. Observations based on the assumption of ideal responses (e.g., Western blotting, or RT-PCR, etc.) may overestimate these peripheral states. BECC can help avoid this risk.

### BECC description and recognitive hierarchy with results obtained in Wet’s experimental system: flow cytometry analysis of phosphorylated Signal Transducer and Activator of Transcription 3 (pSTAT3) using human peripheral blood

An example of BECC is presented using flow cytometry results. The flow cytometry measurements do not represent the ideal responses of single cells (see Confirmation 4) but rather yield results that can be treated as data from a cell population. An additional advantage is the ability to simultaneously analyze multiple parameters, which allows the observer to redefine the target cell population after the measurement using any desired parameter.

Here, we compare BECC across three cell populations: i) BECC based on “white blood cells” defined by cell size and density, ii) BECC based on “T cells” identified using cluster of differentiation (CD) 3 as a marker, and iii) BECC based on “neutrophils” identified using CD16 as a marker. These gating are represented in Fig. 6a. From an immunological perspective, simply using CD3 as a marker does not distinguish among the various T cell subsets, which continue to increase as new subsets are reported; (Zielinski 2023). Similarly, numerous subsets of neutrophils have been identified; (Palomino-Segura et al. 2023). Determining at what level of separation the cell population can be considered homogeneous in terms of response is difficult and varies depending on the experimental conditions. Therefore, we are unable to draw a definitive conclusion. In other words, observation is impossible without first committing to a specific cellular fraction, a decision that precedes the selection of an appropriate one. This suggests that *a priori* classification is standard practice. This state is emulated by the BECC of white blood cells, while the BECCs of CD3⁺ T cells and CD16⁺ neutrophils represent cases where cell populations with a direct association to ideal one-dimensional parameters or experimental conditions can be extracted from white blood cells.

**Fig. 6.**
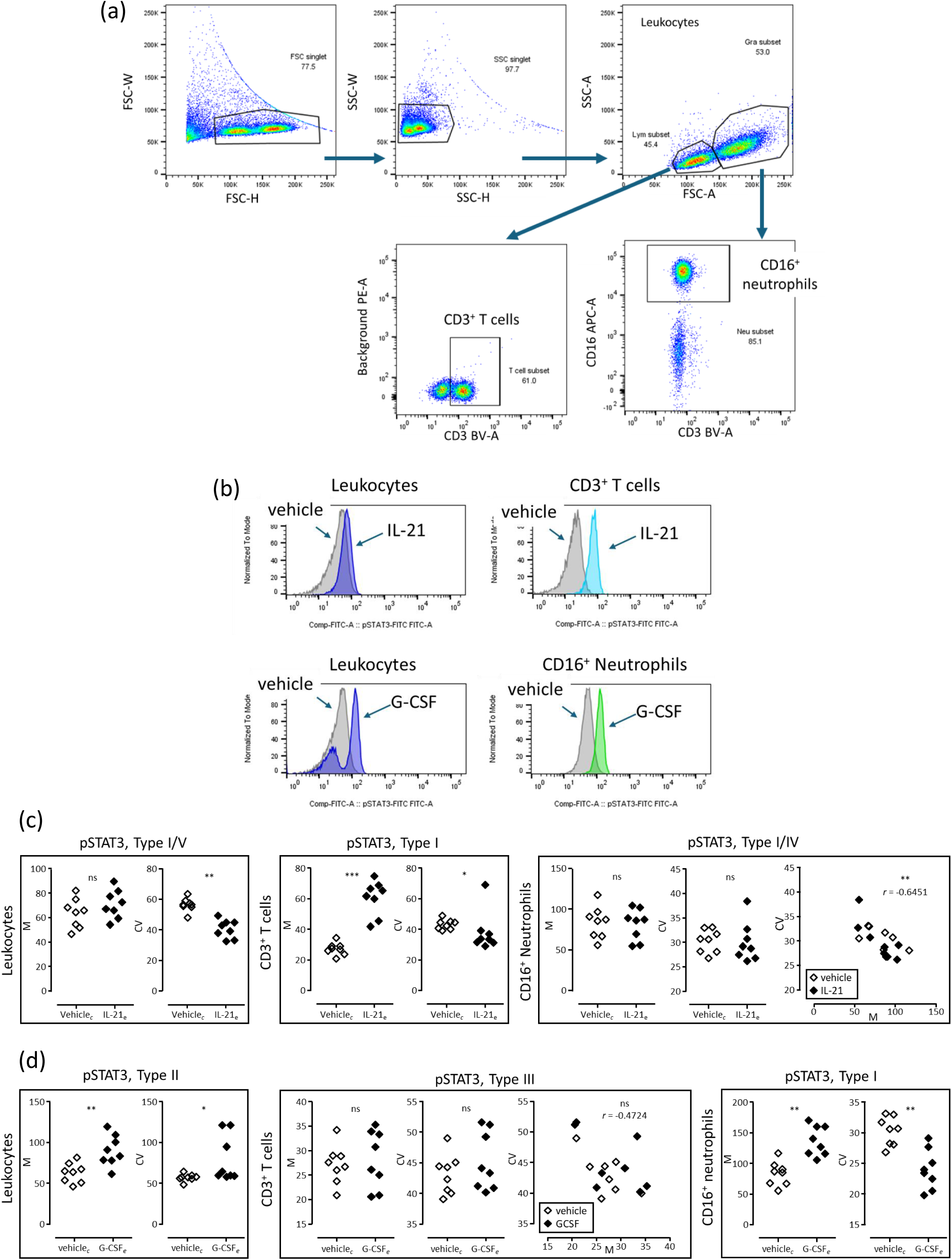
BECC based on STAT3 phosphorylation from flow cytometry. (a) Example of cell fractionation analysis. A representative example is shown of the leukocyte fraction obtained after gating to exclude doublets, and of the granulocyte and lymphocyte fractions. Additionally, a representative example of the analysis of the CD3^+^ lymphocyte (CD3^+^ T cell) and CD16^+^ neutrophil fractions is presented. (b) Representative histogram of phosphorylated STAT3 (pSTAT3) measurement results. The leukocyte fractions 15 minutes after whole blood was treated with vehicle, IL-21 (10 nM), or G-CSF (10 nM) are shown. Vehicle-stimulated leukocyte fractions are indicated in gray, those stimulated with IL-21 or G-CSF are shown in purple, CD3^+^ T cells in light blue, and CD16^+^ neutrophils in green. (c) BECC based on STAT3 phosphorylation results 15 minutes after IL-21 stimulation. The control group (*c*) was treated with vehicle, and the experimental group (*e*) with IL-21. (d) BECC based on STAT3 phosphorylation results 15 minutes after G-CSF stimulation. The control group (*c*) was treated with vehicles, and the experimental group (*e*) with G-CSF. (C and D) BECC results. The rectangles display the combinations of M (mean: mean fluorescence intensity of pSTAT3) and CV (robust coefficient of variation: CV of fluorescence intensity) used in BECC. The left vertical labels of the rectangles indicate cell populations defined by the observer. The top horizontal labels indicate the item measured (pSTAT3) as an ideal one-dimensional variable, and the classification result by BECC. Comparisons between two groups were analyzed using Mann–Whitney U test, and correlation coefficients were analyzed using Spearman’s rank correlation coefficient. ns: not significant; *, *p* < 0.05; **, *p* < 0.01; ***, *p* < 0.001 (*c*, n = 8; *e*, n = 8).

In other point of view, both interleukin (IL)-21 and Granulocyte Colony-Stimulating Factor (G-CSF) are well known to induce the phosphorylation of STAT3 (pSTAT3) as JAK-STAT pathway (Lv et al. 2024). Of course, since each cytokine has a specific receptor, the cells they stimulate are limited. However, when the relationship between a these cytokine receptors and cell population is unknown, an experiment based on an “ideal response” (see Confirmation 4) can only yield the observation of an increase in pSTAT3. Therefore, we conducted an analysis to see how the response could be classified using BECC.

In biological phenomena we aim to observe, heterogeneous cells may already be present. Sometimes they can be separated and observed, while in other cases their existence is not even recognized or anticipated. To demonstrate that, in actual experimental systems, BECC is capable of describing such previously unrecognized heterogeneity, we used leukocytes from human peripheral blood as the cell population and measured pSTAT3 as the ideal one-dimensional parameter. The representative results are shown in Fig. 6b.

At first, we examined the IL21 stimulation. The observation point was set at 15 minutes after stimulation, and we compared the control group *c* (vehicle) with the experimental group *e* (IL-21 stimulation). At 15 minutes, the leukocyte population did not show a significant response in M (the mean of pSTAT3 fluorescence intensity in the cell population) to IL-21 stimulation, but CV (the CV of the pSTAT3 fluorescence intensity on each cell in the cell population) significantly decreased (Fig. 6c, left panel). In terms of BECC, this corresponds to Type I/V. The leukocyte population shows the presence of an unrecognizable pSTAT3 response involved in attraction in response to IL-21.

Focusing on CD3^+^ T cells contained within the leukocyte population as the target cell population, we analyzed pSTAT3. At 15 minutes, the CD3^+^ T cell population showed a significant increase in the pSTAT3 response, and the CV significantly decreased (Fig. 6c, center panel). According to the BECC, this corresponds to Type I. This indicates that the CD3^+^ T cell population is responsive to IL-21 stimulation, with an attract increase in pSTAT3.

Focusing on CD16^+^ neutrophils contained within the leukocyte population as the target cell population, we analyzed pSTAT3. Although there was no significant difference in either M or CV of the pSTAT3, an inverse correlation between M and C was observed (Fig. 6c, right panel). BECC indicates Type I/IV. This suggests that the CD16^+^ neutrophils were induced to pSTAT3 by conditions of non-recognition, regardless of whether they are in the control or experimental group.

The leukocyte population is composed of various cell types, including CD3^+^ T cells and CD16^+^ neutrophils. The overall response observed in the leukocyte population is therefore a result of the combined responses of both CD3^+^ T cells and CD16^+^ neutrophils.

Paradoxically, when we were unable to separate T cells and neutrophils from the leukocyte population, the BECC in response to IL-21 stimulation was classified as Type I/V. While this initially appears to show no clear response, it suggests the possibility of an unrecognized cell population responding to IL-21. Type I/V findings indicate that the presence of unrecognized inducible responses inherent within the cell population.

Next, we examined the G-CSF stimulation. The observation point was also set at 15 minutes after stimulation, and the control group *c* (unstimulated vehicle) was compared with the experimental group *e* (G-CSF stimulation) (Fig. 6b, left panel). As a result, at 15 minutes post-stimulation, both M and CV of pSTAT3 in the leukocyte population significantly increased in response to G-CSF. According to BECC, this corresponds to Type II. Thus, it indicates that the leukocyte population shows a pSTAT3 response that involves the emergence of heterogeneous cells in response to G-CSF stimulation.

Focusing on pSTAT3 in the CD3^+^ T cell population contained within the leukocyte population, the CD3^+^ T cells at 15 minutes post-G-CSF stimulation showed no significant change in either M or CV of pSTAT3, and no correlation was observed (Fig. 6b, center panel). According to BECC, this corresponds to Type III. This suggests that the CD3^+^ T cell population does not respond to G-CSF (i.e., acts as a bystander).

Focusing on the CD16^+^ neutrophils within the leukocyte population, at 15 minutes post-G-CSF stimulation, the CD16^+^ neutrophil population showed an increase in M and a decrease in CV. According to BECC, this corresponds to Type I (Fig. 6b, right panel). This indicates that the CD16^+^ neutrophil population responds to G-CSF stimulation with an inducible pSTAT3 response.

In summary, as in the previous analysis, the leukocyte population includes both CD3^+^ T cells and CD16^+^ neutrophils. The BECC of the leukocyte population in response to G-CSF is classified as Type II, indicating the emergence of heterogeneous cells in response to G-CSF.

We propose the following method to describe BECC in a linear, sentence-like format instead of using graphical panels, such as those in Fig. 6. First, the cell population observed by the researcher is described, followed by the group comparison (control (*c*) and experimental (*e*)), enclosed in square brackets. Finally, the result of the BECC is indicated as a superscript to the ideal one-dimensional marker, incorporating the observer’s perspective into the BECC notation. Specifically:

Fig. 6c left panel: Leukocyte [vehicle - IL-21] pSTAT3ᴵ/ⱽ

Fig. 6d left panel: Leukocyte [vehicle - G-CSF] pSTAT3ᴵᴵ

This notation indicates that pSTAT3 exists in multiple states within the cell population rather than a single uniform state. By focusing on the differences and shared BECC distinctions between cell types, control groups, and experimental groups, a further level of cognition, namely cognitive hierarchy, can be achieved.

An example of this sentence-based description would be: “We focused on STAT3 phosphorylation 15 minutes after cell stimulation for the purpose of analysis. As a result, we observed leukocyte [vehicle - IL-21] pSTAT3ᴵ/ⱽ, whereas leukocyte [vehicle - G-CSF] pSTAT3ᴵᴵ. Based on this, the differing pSTAT3 results from IL-21 and G-CSF stimulation suggest the existence of an unidentifiable event in the relationship between the stimulus and the leukocytes. This unidentifiable event is presumed to be related to a background in which pSTAT3 is induced, and the emergence of a heterogeneous cell population within the leukocytes upon G-CSF stimulation.”

This example serves only as one possible interpretation, as the purpose and understanding vary depending on the observer. The key point to note here is that comparing BECCs allows for the formal realization of cognitive hierarchization. Consequently, this offers a means to surface previously unrecognized events and new perspectives for understanding.

### BECC description and recognitive hierarchy with results obtained in Wet’s experimental system: BECC in T Cells of Ovalbumin-specific TCR transgenic, class II–restricted mice (OT-II) Mice

Next, using OT-II and wild type (WT) mice, we demonstrate that BECC can be applied not only to in vitro molecular reactions or intracellular molecular events but also to more complex, multilayered biological phenomena that occur *in vivo*. OT-II are transgenic mice whose T cells express a T-cell receptor (TCR) that specifically recognizes the ovalbumin (OVA_323–339_) peptide presented on major histocompatibility complex (MHC) class II molecules. This TCR is composed of the TCRα and TCRβ chains, specifically TCR Vα2 (TRAV14) and TCR Vβ5 (TRBV5) (Barnden et al. 1998). The background of OT-II and the relationship between T cell differentiation and T cell receptors (TCR) are shown in Fig. 7a.

**Fig. 7.**
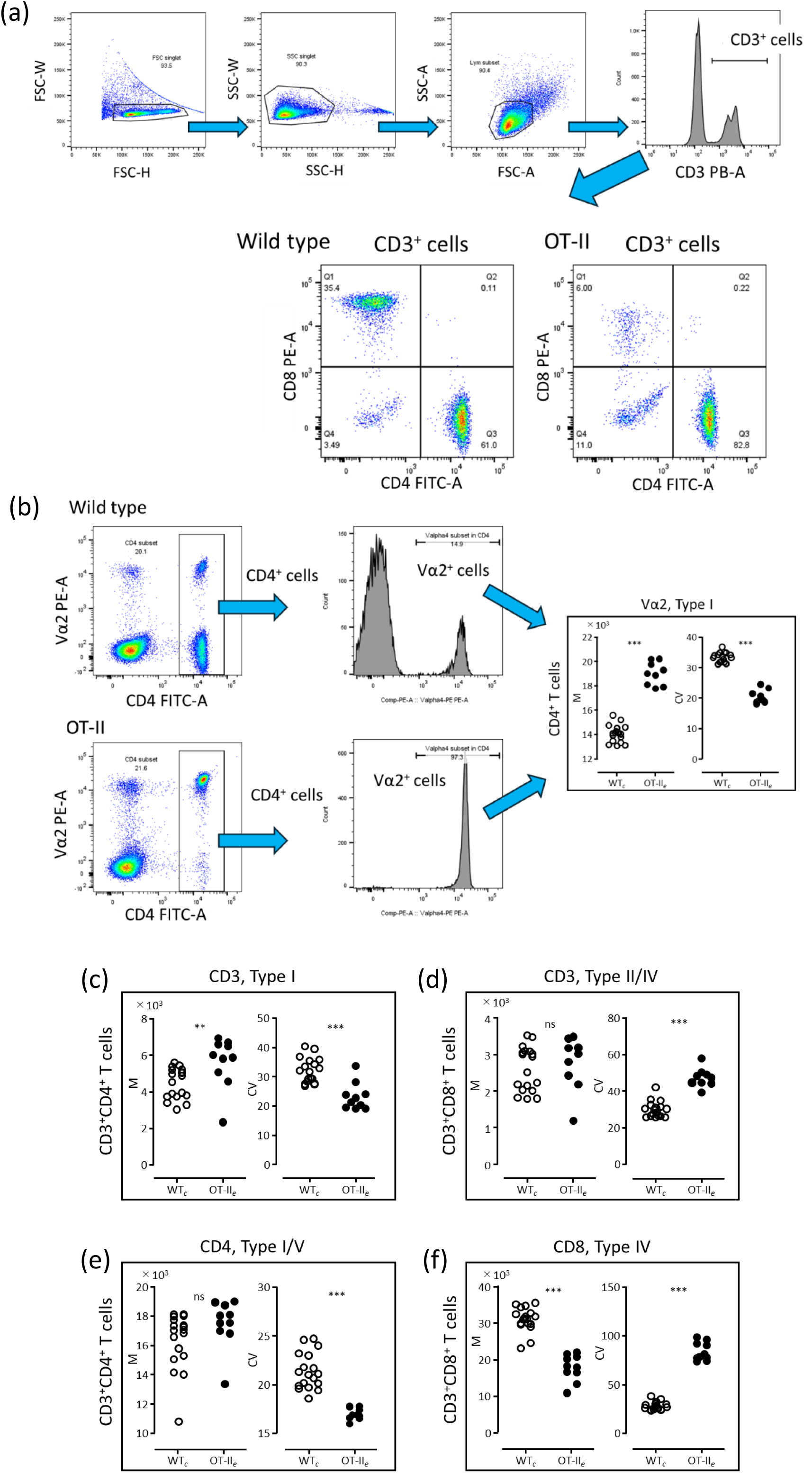
BECC of T Cells in OT-II Mice Overexpressing Vα2 from flow cytometry. (a) Representative analysis of CD4^+^ and CD8^+^ T cells in wild-type (WT) mice and OT-II mice (OT-II). Peripheral blood leukocytes were stained with BV-conjugated anti-CD3 mAb, FITC-conjugated anti-CD4 mAb, and PE-conjugated anti-CD8 mAb. Single cells were selected using gating based on FSC-W,-H and SSC-W,-H, followed by gating of CD3^+^ cells. CD4^+^ and CD8^+^ T cells within the CD3^+^ T cell population were further analyzed. (b) Representative example of Vα2 overexpression in WT and OT-II. Peripheral blood leukocytes were stained with FITC-conjugated anti-CD4 mAb and PE-conjugated anti-Vα2 mAb. Vα2 expression in CD4^+^ peripheral blood leukocytes was analyzed, and BECC was performed by comparing the expression levels of Vα2 between WT and OT-II mice. Statistical analysis was conducted using the Mann–Whitney U test. ***, *p* < 0.001 (*c*: WT, n = 18; *e*: OT-II, n = 10). (C – F) BECC of T cells in WT and OT-II. BECC was performed on the following parameters: CD3 expression in CD4^+^ cells (c), CD3 expression in CD8^+^ cells (d), CD4 expression in CD4^+^ cells (e), and CD8 expression in CD8^+^ cells (f). Statistical analysis was conducted using Mann–Whitney U test. ns: not significant; **, *p* < 0.01; ***, *p* < 0.001 (*c*: WT, n = 18; *e*: OT-II, n = 10).

It is known that during thymic T-cell lineage commitment, T cells differentiate into CD8⁺ cells when their TCRs recognize MHC class I, and into CD4⁺ cells when they recognize MHC class II. Accordingly, T cells in OT-II mice are typically directed toward the CD4⁺ T cell lineage. However, even in such TCRα/TCRβ transgenic mice, endogenous TCR gene rearrangement is not always fully suppressed, and thus a small population of CD8⁺ T cells may still be present (Fig. 7a). These CD8⁺ T cells are expected to possess a diverse repertoire of TCRs unrelated to the transgenically overexpressed TCR gene segments (TRAV14 and TRBV5). We applied BECC to describe T cells derived from OT-II mice by comparing them with normally differentiated T cells from WT mice.

In OT-II T cells, TCRs containing gene fragments derived from the Vα2 gene are overexpressed and compared to wild-type (WT) mice. Compared with WT mice, T cell differentiation is directed toward Vα2-positive CD4⁺ T cells. Therefore, we examined whether the description by BECC aligns with this genetically induced bias (Fig. 7b). The cell population was defined as CD4⁺ cells, and the expression level of Vα2 was used as the ideal one-dimensional feature. The control group (*c*) was WT, and the experimental group (*e*) was OT-II. As a result, the BECC of CD4⁺ cells in OT-II were identified as Type I. This indicates that Vα2 expression is indeed induced in OT-II. As shown in Fig. 7b, the flow cytometry results were able to describe the Vα4 gene overexpression operation in OT-II as Type I (M↑, CV↓; Attraction). Accordingly, the CD4^+^ T cells that overexpress Vα4 in OT-II are presented as

Fig. 7b: CD3^+^CD4^+^ [WT–OT-II] Vα4^I^ using BECC.

The surface expression of the CD3 complex, which acts as the signaling unit, reflects cooperative behavior and is considered a molecule associated with the overexpressed Vα4 components. The TCR complex consists of antigen-recognition units, TCRα and TCRβ chains, and the signaling unit, known as the CD3 complex, which includes CD3γ, CD3δ, CD3ε, and ζ chains. In OT-II mice, TCRα and TCRβ chains are overexpressed, whereas the expression of the CD3ε chain is derived from the endogenous genetic background. At the protein level, the TCRαβ heterodimer and CD3 complex chains associate appropriately within the cell, become transported to the cell surface, and form a stable, functional TCR complex on the membrane. Therefore, surface expression of the CD3 complex, which acts as the signaling unit, reflects cooperative behavior and is considered a molecule associated with the overexpressed Vα4 components. We thus examined how the CD3 complex is represented by BECC when the antigen recognition unit of the TCR is induced in OT-II mice (see Fig. 7c and 7d).

The cell populations were defined as either CD4⁺ cells (Fig. 7c) or CD8⁺ cells (Fig. 7d), with the CD3ε expression level used as the ideal one-dimensional feature. As a result, BECC in CD4⁺ cells of OT-II were classified as Type I. In other words, this Type I result indicates that functional coordination of TCRs recognizing MHC class II was being induced in OT-II. On the other hand, in CD8⁺ cells of OT-II, BECC was classified as Type II/IV. This reflects the presence of CD8⁺ T cells that differentiated due to incomplete suppression of diverse endogenous TCR gene rearrangements. Taken together, these results demonstrate that BECC can categorize associated peripheral biological phenomena, i.e., multistage and multilayered events, arising from genetically induced changes.

The mechanisms of T cell differentiation are well known (Taniuchi 2018). During T-cell differentiation, CD4 and CD8 expressions begin when bone marrow-derived lymphoid progenitor cells migrate to the thymus and become Double Negative (DN) cells, which do not express either CD4 or CD8. These DN cells then undergo TCR gene rearrangement. Upon successful expression of a functional TCRβ chain, the cells differentiate into Double Positive (DP) cells expressing both CD4 and CD8. If the TCR of a DP cell binds to MHC class II, intracellular signaling pathways are activated to maintain the expression of the CD4 gene while suppressing CD8 gene transcription, leading to differentiation into CD4 Single Positive (SP) cells. Conversely, if the TCR recognizes MHC class I, the cell maintains CD8 gene expression while suppressing CD4 transcription, differentiating into CD8 SP cells. Therefore, in OT-II mice, where TCR genes that recognize MHC class II are overexpressed, differentiation into CD8 SP cells, and thus CD8 expression, is indirectly suppressed.

In this study, peripheral blood was used for measurements, meaning the cells analyzed were mature SP cells that had completed the differentiation control processes. Initially, we assumed that in peripheral blood, where these processes had ended, there would be no difference between WT and OT-II mice in terms of CD4 and CD8 expression levels or their CV. However, unexpectedly, BECC of CD4 expression in CD4⁺ T cells of OT-II mice were classified as Type I/V (Fig. 7e), and BECC of CD8 expression in CD8⁺ T cells were classified as Type IV (Fig. 7f). This indicates that the expression of CD8, an indirectly associated event with the overexpressed TCR, can indeed be observed as BECC. The mechanisms that induce T-cell differentiation are well understood at the molecular level and were used to create the OT-II mouse model. To explain how BECC can be used to handle unrecognized events, we will now reinterpret the OT-II mouse not as a genetically engineered model, but rather as a mutant discovered by chance, and assume that the molecular mechanisms of T-cell differentiation are also unknown.

In this hypothetical framework, we state: Compared to WT, we identified a mutant mouse, OT-II, with an increased proportion of CD4⁺ T cells. The cause was unknown.

The results are able to describe as follows: “The observer compared WT and OT-II in terms of CD3 expression, CD4 expression, and CD8 expression, using CD4⁺ and CD8⁺ T cells as cell populations and the markers as ideal one-dimensional features. As a result, in OT-II, CD3 expression in CD4⁺ T cells were classified as Type I, and CD4 expression was classified as Type I/V. In CD8⁺ T cells, CD3 expression was classified as Type II/IV, while CD8 expression was classified as Type IV. Thus, in OT-II, CD3 expressions in CD4⁺ T cells were increased and appeared to be induced, while CD8 expression in CD8⁺ T cells were decreased and suggested the emergence of heterogeneous cells. CD4 expression in CD4⁺ T cells of OT-II might have been induced, and CD3 expression in CD8⁺ T cells might have indicated the emergence of heterogeneous populations.”

We then use linear notation for BECC instead of graphical panels. In this case, since the cell population indicators and the ideal one-dimensional features were the same, the BECC notation could be written as:

Fig. 7c and 7e: CD3^+I^ CD4^+I/V^ [WT–OT-II]) Fig. 7d and 7f: CD3^+II/IV^ CD8^+IV^ [WT–OT-II]

In BECC, odd-numbered types (excluding Type III) indicate induction, while even-numbered types point to the emergence of heterogeneous cells. The elements involved in induction are useful for causally interpreting the differences between the WT and OT-II mice. Conversely, the elements showing heterogeneity help us interpret the phenomena associated with these differences. Therefore, our causal interpretation of this phenomenon lead us to focus on CD3⁺ᴵ. At the same time, we pay attention to CD8^+IV^ as a symmetrically associated phenomenon.

Ultimately, BECC is a valuable tool for recognition of biological phenomena. It helps to clearly define the boundary of the unknown, which enables the discovery of novel phenomena.

## Discussion

At the core of approaches to understanding biological phenomena lies the problem of the observer—their existence and inherent arbitrariness. In this study, we explicitly incorporated the observer into our framework, enabling us to define minimal unit of recognition and life, and to clarify the relationship between time and experimental error in observation. Building on these assumptions, we classified cellular behavior into 11 distinct patterns, validated through agent-based modeling simulations.

As a practical demonstration, we applied this framework to intracellular signaling and cell differentiation, using flow cytometry data. As a result, BECC demonstrated its ability to describe biological phenomena, including unidentified cells and undiscovered events, prior to the recognition of empirical facts. This resolves the problem of reproducibility, which is a major obstacle to open science in life sciences. Furthermore, it provides a means to solve issues related to linguistic fluctuations and conservative recognition as below.

The issue of reproducibility arises from the inability to identify differences that are inherently present in an unrecognizable form, as well as the difficulty of integrating numerical data. The increase in CV observed in BECC, expresses these “inherently unrecognizable differences” as the presence of unsynchronized “heterogeneous cells” within the ideal one-dimensional parameter. Furthermore, in BECC, once the data are classified into 11 distinct types, they are liberated from numerical constraints. For example, regardless of when or where the measurement is taken, the result can be expressed as “cell behavior classified as Type I.” This approach helps resolve reproducibility issues. It also allows us to suppress the overestimation of experimental results that, unlike BECC, assume ideal reactions (see Confirmation 4).

Regarding the issue of linguistic fluctuation, BECC allows for the expression of “different recognitions” that are independent of linguistic representations. Type I is not a “strong Type I” or a “typical Type I”; instead, it is a symbol that conveys only the behavior of Type I cells. The BECC classification notation then separates recognition of results from understanding. Even if an observer overstates the understanding of a small-scale experiment, the BECC results remain unchanged. In other words, focusing on BECC allows one to isolate the “spin” of a scientific paper. Thus, the results are not subject to error across different research domains, and inflated interpretations or overstated concerns can be readily distinguished.

Moreover, because BECC represents the minimal unit of recognition derived from an ideal one-dimensional parameter, classifications based on BECC cannot be further subdivided. In other words, BECC classification delineates the boundaries of recognition that arise from the arbitrariness of the observer. Consequently, it avoids descriptions that rely on implicitly assumed “unrecognized differences” within disciplines (for example, explaining specific signaling pathways in terms of universal pathways, such as assuming that a reaction is specific merely because it is inhibited by the same inhibitor that suppresses another reaction). As a result, misinterpretations across different fields can be prevented.

The issue of conservative recognition can likewise be addressed through BECC. Observers are required to make an explicit “declaration”—that is, to present the experimental conditions defining the control (*c*) and experimental (*e*) groups—based on an ideal one-dimensional parameter for BECC classification. Consequently, if the observed results yield a classification type that differs from previous studies, it becomes evident whether the discrepancy arises from the observer’s declaration or from an inherently unrecognizable difference. In other words, even when adherence to prior declarations reflects conservatism, BECC-based recognition remains readily adaptable to new observations. By interrogating the reasons underlying BECC differences, researchers can move beyond conservative modes of understanding. For instance, BECC enables recognition that is not constrained by dualistic frameworks such as inflammation versus anti-inflammation or promotion versus inhibition, thereby facilitating a more advanced understanding of biological phenomena.

The classification framework, BECC, provides approach to *a priori* recognizing such non-empirical understanding. BECC is not intended to directly analyze or uncover new mechanisms or phenomena. Rather, it serves as a descriptive framework designed to address issues of reproducibility, linguistic fluctuation, and conservative recognition. Consequently, it enables the destruction and sharing of boundaries that would otherwise remain unrecognized. This, in turn, facilitates the identification of novel phenomena through shared recognition of patterns previously overlooked.

Variability in flow cytometry instruments leads to differences in M and CV, as do the staining properties of fluorescent dyes and variations in detection sensitivity. Similarly, comparisons based on the percentage of positive cells may yield divergent results— observations that are familiar to many researchers. However, the patterns of increase or decrease in M and CV observed between two groups constitute a “structure” that remains consistent regardless of the measuring instrument or its sensitivity. In other words, BECC can be applied by simply comparing M and CV across two groups, without requiring global standardization. This approach not only mitigates problems related to the reproducibility of primary data but also enhances the reliability of open science within the life sciences. Ultimately, BECC functions as a method for classifying “structures of difference” rather than absolute “values.”

Analyzing cell populations without assuming ideal responses is straightforward when employing flow cytometry. However, this does not imply that flow cytometry is the only experimental tool suitable for BECC. The definition of a “cell” is inherently arbitrary and ultimately determined by the observer. A “cell” may be defined at the level of an individual organism, in which case behavioral analyses can also be adapted to BECC. At the molecular level, if individual molecules are regarded as “cells” and analyzed collectively as a population without assuming ideal responses, BECC can likewise be applied to fluctuations in molecular reactions within complex systems.

For BECC to be applied across diverse experimental systems, four conditions must be met: (i) a description of arbitrarily defined “cells”; (ii) a description of ideal one-dimensional parameters; (iii) a description of control and experimental groups; and (iv) a statistical description, as discussed below.

At present, no logically definitive method exists for determining the appropriate number of replicates in statistical analyses. In most cases, the number of experimental replicates is dictated by practical constraints such as feasibility and cost. Within BECC, because significance testing is conducted using the Mann–Whitney test, a minimum of four experimental replicates is statistically required. Furthermore, as demonstrated in simulation studies, CV tends to fluctuate markedly during both the early and late phases of a response. For this reason, we generally performed more than six experimental replicates before conducting statistical analyses.

In routine flow cytometry, performing approximately six replicates is relatively feasible. By contrast, conducting six replicates of mass cytometry or scRNA-seq is often economically impractical. In such circumstances, we do not exclude the possibility that some investigators may consider a parametric t-test, under the assumption of normality, to be sufficient. Nonetheless, it is essential to emphasize that verifying genuine normality requires a substantial number of replicates. Reliance on a presumed normal distribution based on as few as three replicates should instead be regarded as a form of “conservative recognition.”

We now live in an era in which vast numbers of new papers are generated through artificial intelligence (AI)–based summarization of existing literature and the use of publicly available data. Without a clear understanding and shared recognition of how primary data are formed, such AI-generated manuscripts risk becoming little more than efficiently assembled anagrams. As a countermeasure, BECC provides a promising framework. Because its implementation requires satisfaction of the four conditions outlined above, BECC enables recognition to be distinguished from interpretation, thereby improving our understanding of the processes underlying primary data formation. In this way, BECC offers a practical means of validating AI-generated output while maintaining framework of *a priori* recognition.

BECC was a classification method that used the minimal unit for recognizing biological phenomena. By applying this approach, it will become possible to transcend issues of reproducibility, linguistic fluctuation, and conservative recognition. This facilitates the sharing of cognitive boundaries with observers who exist in different spatial and temporal contexts. Therefore, BECC will serve as a descriptive method that promotes open science in life sciences.

## Additional Information

### Author contributions

Y.T. conceived the study. Y.T. performed simulations and flow cytometric experiments for the study. Y.T. wrote the manuscript. J.Y., R.Y., S.S., and H.A. reviewed the manuscript.

### Ethics

Human blood collection: This study was approved by the Ethics Committee of Yamagata University Faculty of Medicine (approval number: 2024-44).

Mice blood collection: The animal experiments were approved by the Animal Experiment Committee of Yamagata University Faculty of Medicine (approval number, R7015).

## Additional files

### Supplementary files

Supplementary Information

## Data availability

The NetLogo file (Model of cell population_ver.5.02) has been deposited in Zenodo (DOI: 10.5281/zenodo.17230524).

All raw data have been deposited in Zendo (DOI: 10.5281/zenodo.17232980).

