## Supplementary Information for "*A Priori* recognition for biological phenomena prior to empirical understanding"

**Supplementary Information: About the NetLogo Simulation Program "Model of Cell Population_ver.5.0.2"**

**1. Fixed parameter values**

To make simulations easily executable on personal computers used by observers, limitations are placed on the size of the *World* (the interface screen visualizing agent behavior; see Figure 1B). A setup that requires lengthy computations for the simulation to complete would not allow for easy verification. While the size of the *World* and other settings can be changed via the *interface tab*, as these settings affect visualization, users may adjust them based on their computer environment.

As a recommended setup, we configured the *World* size and agent behavior so that the simulation concludes within approximately 5,000 to 10,000 ticks. Specifically, the *World* size was set to **201 × 201**, with a patch size of **3**, a frame rate of **10**, and looping enabled in all directions. The **activator** moves **0.3 units per tick** (randomly turning left or right within a 0–40° range), while the **cell** also moves **0.3 units per tick** (with random left-right movement within a 0–360° range).

Although such fixed settings may affect the timing of the overall agent response, they do not act as parameters that artificially influence the measured M or CV values in the simulations of this study.

The behavior of agents as represented in the *World* is useful for verifying the program code and for intuitive understanding. Therefore, it's important to make agent behavior visually accessible for human observers. To that end, the visualization was designed as follows:

- The target cells (Target-cell) are shown in cyan, shaped as circles, and sized 3.
- Heterogeneous cells (Hetero-cell) are assigned random colors, excluding cyan, white, red, yellow, and No.18 (light red), to make them easily distinguishable. Their shape and size match those of the Target-cells: circle and 3.
- The activators are represented in red, shaped as triangles, and sized 2.
- The patch color of the background is set to No.18, a tone chosen for clear contrast with both cells and activators.
- Once a cell begins responding, it changes to yellow, with the shade of yellow varying depending on its response value.

These agent behaviors are described in more detail below. No behavioral instructions are assigned to the background patches apart from color designation.

**2. Behavior of the Activator**

The activator serves as the conceptual trigger for a reaction—meaning that a reaction begins when an activator encounters a cell. Once an activator and a cell have met, the arrival of an additional activator or cell at the same location does not initiate a new reaction trigger. Furthermore, after serving as the reaction trigger, the activator disappears. Activators appear at random positions within the *World*. If too many activators are present, they may encounter cells instantly, which would cause the number of activators to artificially dictate the synchrony of reactions. To prevent this, the simulation is configured so that only one activator appears per tick. This behavior allows for focused examination of the relationship between the frequency of activator–cell encounters (reaction triggers) and CV.

**3. Behavior of the Target Cell**

The NetLogo file used in this study, *"Cell Population Model ver. 5.0.2,"* allows each individual cell to be assigned random values for the following parameters: “initial response value (held at the time of appearance),” “rate of increase in response value,” “duration of maintained response,” and “rate of decrease in response value” (see “Adjustable Parameter Values: Slider Descriptions” below). However, if these values are completely unrestricted, the result would be total disorder.

In this context, the cells selected by the observer as Target-cells must be similar to each other. This means that the range of randomness assigned to them is constrained to a certain extent, in order to represent similarity among cells.

Furthermore, once a cell’s reaction ends—specifically, when its response value falls below the “initial response value”—it becomes capable of reacting again. This behavior allows for focused analysis of the relationship between how the Target-cell responds and the resulting CV (coefficient of variation).

**4. Behavior of the Hetero-cell**

The Hetero-cell represents pre-existing heterogeneous cells. When the proportion of Target-cells is reduced, Hetero-cells appear in the *World* corresponding to the reduced ratio. Whether Hetero-cells react upon encountering an activator is determined by the switch **"Hetero-cells-response-to-act"**, which can be toggled **ON** or **OFF**.

If set to **OFF**, Hetero-cells do not react even when they encounter an activator (and the activator does not disappear). If set to **ON**, Hetero-cells can react upon encountering an activator in the same way as Target-cells. However, unlike Target-cells, each Hetero-cell is independently assigned random values for the **rate of response increase**, **duration of maintained response**, and **rate of response decrease**—all within their own random ranges. As a result, Hetero-cells exhibit different response behaviors from Target-cells.

This behavior allows for focused examination of the relationship between the presence of heterogeneous cells (Hetero-cells) within the Target-cell population and the resulting CV (coefficient of variation).

**5. Adjustable Parameter Values**

The *“Cell Population Model ver.5.0.2”* includes several parameters that can be adjusted before starting the simulation, much like setting experimental conditions. These parameters are controlled using sliders located in the *interface tab* (Fig. 2B). If you wish to change a value beyond the preset range of a slider, and your operating system is Windows, you can right-click on the slider, select “Edit” from the pop-up window, and freely redefine the range of values. Similarly, when performing a parameter sweep (via *Tools → BehaviorSpace → Edit*), you can manually input any desired values. The adjustable parameters are described below.

***5.1. Sliders on the right side of the screen***

- **Total-cell-num**: Adjusts the total number of cells that appear in the *World*. A range of approximately 1,000 to 2,000 cells is appropriate for visual observation within the *World*.
- **Target-cell-%**: Changes the proportion of cells, relative to the total cell count, that the observer considers to be the target cells. To simulate the impact of heterogeneous cells, reduce this percentage.
- **Finish-ticks**: Sets the number of ticks at which the simulation ends. A range of around 5,000 to 10,000 ticks provides a suitable waiting time for visual observation in the *World*.
- **Activator-num**: Adjusts the number of activators that appear in the *World*. The number should be set relative to the total number of cells; values near this baseline are optimal for visual tracking. To increase the frequency of reactions, raise this number.

***5.2. Sliders in the Center of the Screen***

Below the label **“Target-cell’s random-range,”** two columns of sliders are arranged. These sliders allow the user to set the range of randomness (as noise) for the following properties of the Target-cells, as defined by the observer: initial response value, rate of increase, duration of maintained response, and rate of decrease.

- **basal-response** and **basal-noise**: These control the initial response value and the range of randomness applied to it. Increasing the value of basal-noise raises the CV of the initial values. The initial response value assigned to each Target-cell serves as the minimum threshold of that cell's response value.
- **act-rate and act-rate-noise**: These adjust the rate at which the response value increases during the active response period, and the range of randomness applied to that rate.
- **deact-rate** and **deact-rate-noise**: These adjust the rate at which the response value decreases after the response period ends, and the range of randomness applied to that rate. The response value continues to decrease until it reaches each cell’s minimum threshold (its initial response value).
- **retention** and **retention-noise**: These set the number of ticks during which the cell remains in the reactive state, and the range of randomness for that duration. After a Target-cell encounters an activator and the specified number of ticks has passed, the reaction ends and the response value begins to decrease according to the set rate.

***5.3. Sliders on the Left Side of the Screen***

Below the label **“Hetero-cell’s random-range,”** two columns of sliders are displayed. These sliders allow you to independently set the range of randomness (as noise) for the following parameters of non-target cells (Hetero-cells): rate of increase, duration of maintained response, and rate of decrease. Each parameter is prefixed with **“hetero-”** to distinguish it from the variables used for Target-cells. Since the meaning of each variable for Hetero-cells is the same as that for Target-cells, further explanation is omitted here.

**6. “Setup” and “Go/Stop”**

Before starting the simulation, press the **“setup”** button to initialize the *World*. This resets the tick counter to its initial value and generates each agent on the patches and turtles according to the specified conditions. After that, the simulation is started using the **“go/stop”** button. If you wish to pause or restart the simulation, press the **go/stop** button again.

**7. Monitors (Numerical displays)**

In the middle section of the *Interface tab*, there are six horizontally aligned monitors. From left to right on the screen, they are as follows:

- **Hetero cells** and **Responding hetero-cells**: Display the total number of generated Hetero-cells and the number of Hetero-cells currently responding during the simulation.
- **Target cells** and **Responding target cells**: Display the total number of generated Target-cells and the number of Target-cells currently responding.
- **Total cell number** and **Total responding cells**: Display the total number of all generated cells and the number of all cells currently responding during the simulation.

**8. Plots**

At the bottom of the *Interface tab*, there are four plotting windows arranged in a 2x2 layout: **CV**, **Histogram**, **Mean**, and **Activator**. After the simulation begins, the following data from all cells appearing in the *World* are plotted:

- **CV**, **Mean**, and **Activator**: The horizontal axis represents ticks, showing time-series changes.
- **Histogram**: The horizontal axis represents response values, and the vertical axis represents the number of cells. The histogram updates with each tick.

**9. Actions to be performed by the observer**

In Model of cell population file ver. 5.0.2, the observer's action is to press the "**set**" button, and the **"go/stop**" button on the interface screen. Alternatively, to perform a parameter sweep, select "Behavior Space" from the "tool" pull-down menu and click the "**go/stop**" button. The results of the parameter sweep will be output as a table of values in CSV format. The output values are lined up with M and CV calculated for each tick (each agent in the model performs one action based on the instructions of the program code), so the values for any tick can be used for analysis. The entity that performs the above actions is the observer.

**10. Interface tab**

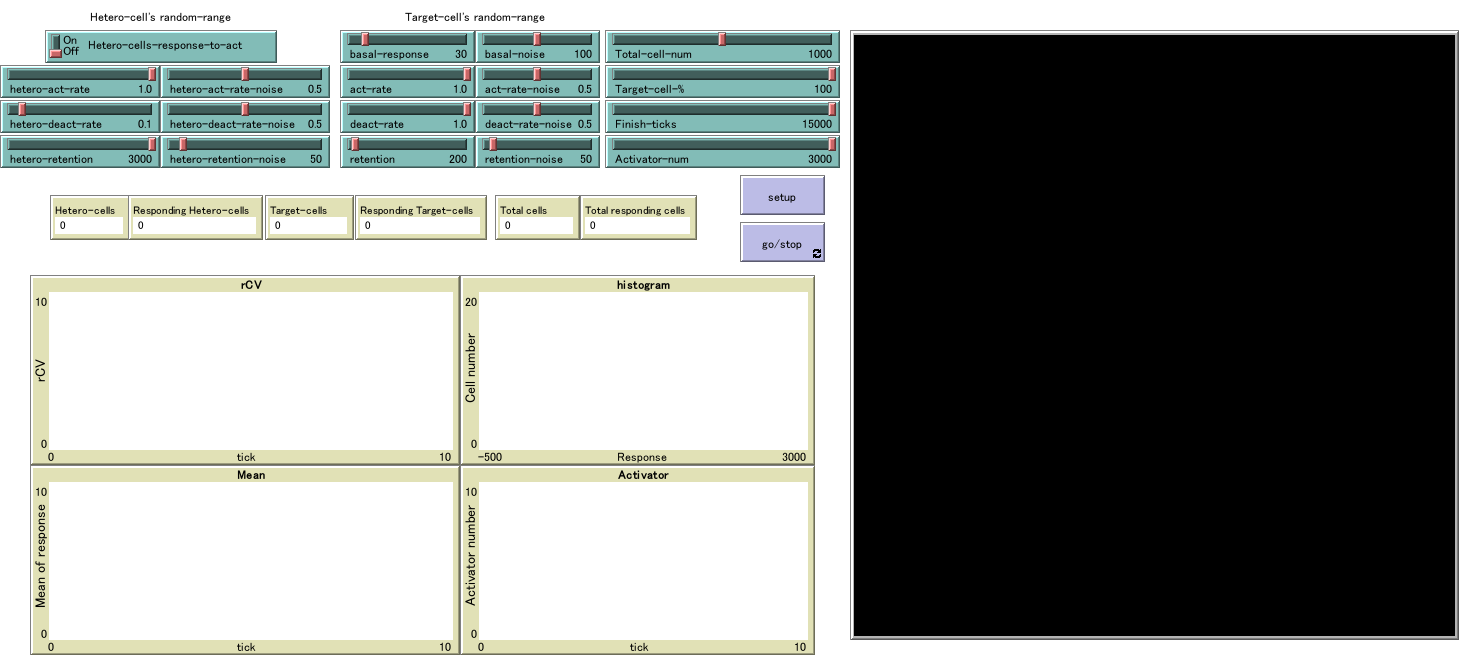

**11. Program code**

breed [activators activator]	; Factors (symbol) that elicit a response

breed [Target-cells Target-cell] ; A cell population the observer wants to measure

breed [Hetero-cells Hetero-cell] ; An unexpected cell population for the observer

turtles-own	;Variables held by each cell

[response	; Value to which the cell responded

responding-time	; Elapsed time (tick) to start response

;The following are values conditioned by the observer

retention-time	; Responding period

activation-rate	; Ratio of increase in response

deactivation-rate	; Ratio of decrease in response

response-lower-limit ; Lower limit of response (= initial value of "response")

B-retention-time

B-activation-rate

B-deactivation-rate]

to setup

clear-all

setup-patches

setup-turtles

reset-ticks

end

to setup-patches

ask patches [set pcolor 18]

end

to setup-turtles ; Cells are appear in the specified number and ratio of cells by the observer.

create-Target-cells (Total-cell-num * Target-cell-% * 0.01) [

set shape "circle"

set size 3

setxy random-xcor random-ycor

set color cyan

set responding-time 0

let r-retention random retention-noise

set retention-time (r-retention + retention)

let r-response random basal-noise

set response (r-response + basal-response)

let r-activation-rate random-float act-rate-noise

set activation-rate (r-activation-rate + act-rate)

let r-deactivation-rate random-float deact-rate-noise

set deactivation-rate (r-deactivation-rate + deact-rate)

set response-lower-limit response ; The initial value of "response" is same as the value of "response-lower-limit" in each cell.

]

create-hetero-cells (Total-cell-num - Total-cell-num * Target-cell-% * 0.01) [

set shape "circle"

set size 3

setxy random-xcor random-ycor

set color random 140

set responding-time 0

let br-retention random hetero-retention-noise

set B-retention-time (br-retention + hetero-retention)

let r-response random basal-noise

set response (r-response + basal-response)

let Br-activation-rate random-float hetero-act-rate-noise

set B-activation-rate (Br-activation-rate + hetero-act-rate)

let Br-deactivation-rate random-float hetero-deact-rate-noise

set B-deactivation-rate (Br-deactivation-rate + hetero-deact-rate)

set response-lower-limit response

randomize] ;Procedures to ensure that the "Hetero-cell" color does not overlap with other turtles and patch for ease of viewing.

end

to randomize

ask Hetero-cells [if

(color = red) or (color = cyan) or (color = white) or

(color = yellow) or (color = 18) [

set color random 140

randomize]]

end

to go

if ticks <= Activator-num

[create-activators 1 [

set color red

set shape "triangle"

set size 2

setxy random-xcor random-ycor

]] ; Generate one "Activotor" per tick up to the specified number ("Actvator-num").

ask activators [fd 0.3

rt random-float 40

lt random-float 40]

ask calculate-total-cells [fd 0.3 rt random 360 lt random 360 ] ; "calculate-total-cells" are evrithing expect "Activator", see below "to-report calculate-total-cells"

ask Target-cells [if count activators-here = 1 [if responding-time = 0

[set responding-time 1

set color scale-color yellow response 10 400]]]

;When "Target-cells" matches "Activator" and the response has not started (responding-time = 0), the response start time is 1.

;At this time, the color changes to yellow for easy viewing. The yellow tint becomes brighter as the value of "response" increases.

if Hetero-cells-response-to-act ; The procedures when "Hetero-cells" chooses to respond to "Activator" using the switch "Hetero-cells-response-to-act".

[ask Hetero-cells [if count activators-here = 1 [if responding-time = 0

[set responding-time 1

set color scale-color yellow response 10 400]]]]

Cell-activation

tick

if ticks = Finish-ticks [stop]

end

to Cell-activation

ask activators

[if (count Target-cells-here with [responding-time <= 1] >= 1)

[die]] ; "activators" disappears once the cell has initiated a response; "activators" are not affected by the "Target-cell" that is responding.

if Hetero-cells-response-to-act

[ask activators

[if(count Hetero-cells-here with [responding-time <= 1] >= 1)

[die]]]

ask Target-cells [if responding-time >= 1

[if responding-time < retention-time

[set response (response + activation-rate)

set responding-time (responding-time + 1)

set color scale-color yellow response 10 400]]]

; The resopnse is elevated in cells where the response has started and the retention time has not elapsed.

ask Target-cells[if responding-time >= retention-time

[if response >= response-lower-limit

[set response (response - deactivation-rate)

set color scale-color yellow response 10 400]]]

; Cells whose response period exceeds the retention time, the response is reduced, to the lower limit of the response.

if Hetero-cells-response-to-act

[ask Hetero-cells [if responding-time >= 1

[if responding-time < B-retention-time

[set response (response + B-activation-rate)

set responding-time (responding-time + 1)

set color scale-color yellow response 10 400]]]]

if Hetero-cells-response-to-act

[ask Hetero-cells[if responding-time >= B-retention-time

[if response >= response-lower-limit

[set response (response - B-deactivation-rate)

set color scale-color yellow response 10 400]]]]

Re-response

end

to Re-response

ask Target-cells

[if count turtles-here = 1

[if 1 <= responding-time

[if response <= response-lower-limit

[set color cyan

set responding-time 0

let r-response random basal-noise

set response (r-response + basal-response)

set response-lower-limit response]]]]

; If the value of "response" in responding cell (responding-time >= 1) drops below the lower limit, it becomes a new cell capable of responding (responding-time = 0).

if Hetero-cells-response-to-act

[ask Hetero-cells

[if count turtles-here = 1

[if 1 <= responding-time

[if response <= response-lower-limit

[set color random 140

set responding-time 0

let r-response random basal-noise

set response (r-response + basal-response)

set response-lower-limit response

randomize]]]]]

end

to-report calculate-total-cells

let total-cells turtles with [color != red]

report total-cells

end

to-report cal-mean

let mean-response mean [response] of calculate-total-cells

report mean-response

end

to-report cal-median

let median-response median [response] of calculate-total-cells

report median-response

end

to-report calculate-p84.13-values ; Percentiles are calculated using an exclusive method.

let cell-num count calculate-total-cells

let raw-p84.13-rank (1 + cell-num) * 0.8413

let integer-p84.13-rank round raw-p84.13-rank

let p84.13-rank integer-p84.13-rank

if integer-p84.13-rank > raw-p84.13-rank [

set p84.13-rank (integer-p84.13-rank - 1)]

let decimal-p84.13-rank (raw-p84.13-rank - p84.13-rank)

let sorted-response-values sort [response] of calculate-total-cells

let front-p84.13-value item (p84.13-rank + 1) sorted-response-values ;In NetLogo, the order ranking starts from 0 (the 0th is the 1st), so it shifts by one.

let hetero-p84.13-value item p84.13-rank sorted-response-values

let p84.13-values front-p84.13-value + (hetero-p84.13-value - front-p84.13-value) * decimal-p84.13-rank

report p84.13-values

end

to-report calculate-p15.87-values

let cell-num count calculate-total-cells

let raw-p15.78-rank (1 + cell-num ) * 0.1578

let integer-p15.78-rank round raw-p15.78-rank

let p15.78-rank integer-p15.78-rank

if integer-p15.78-rank > raw-p15.78-rank [

set p15.78-rank (integer-p15.78-rank - 1)]

let decimal-p15.78-rank (raw-p15.78-rank - p15.78-rank)

let sorted-response-values sort [response] of calculate-total-cells

let front-p15.78-value item (p15.78-rank + 1) sorted-response-values

let hetero-p15.78-value item p15.78-rank sorted-response-values

let p15.78-values front-p15.78-value + (hetero-p15.78-value - front-p15.78-value) * decimal-p15.78-rank

report p15.78-values

end

to-report cal-rCV

let rCV (calculate-p84.13-values - calculate-p15.87-values) / cal-median * 50

report rCV

end

to-report total-responding-cells

let all-responding-cells (count turtles with [responding-time >= 1])

report all-responding-cells

end

to-report responding-cells

let all-target-responding-cells (count Target-cells with [responding-time >= 1])

report all-target-responding-cells

end

to-report hetero-responding-cells

let all-hetero-responding-cells (count Hetero-cells with [responding-time >= 1])

report all-hetero-responding-cells

end
